# Using occipital ⍺-bursts to modulate behaviour in real-time

**DOI:** 10.1101/2022.09.21.508882

**Authors:** Irene Vigué-Guix, Salvador Soto-Faraco

## Abstract

Spontaneous oscillatory neural activity can influence the processing of incoming sensory input and subsequent behavioural reactions. Spontaneous oscillatory activity mostly appears in stochastic bursts, but typical trial-averaged approaches fail to capture this. We aimed at relating oscillatory bursts in the alpha band (8-13 Hz) to behaviour directly, via an EEG-based brain-computer interface (BCI) that allowed for burst-triggered stimulus presentation in real-time in a visual detection task. According to alpha theories, we hypothesised that targets presented during alpha-bursts should lead to slower responses and higher miss rates, whereas targets presented in the absence of bursts should lead to faster responses and higher false alarm rates. Our findings support the role of bursts in alpha-oscillations in visual perception and exemplify how real-time BCI systems can be used as a test bench for brain-behavioural theories.

## INTRODUCTION

Fluctuations in neural excitability, reflected in ongoing brain oscillations, are thought to impact the visual processing of sensory inputs and behavioural outcomes (Benwell et al., 2017; Benwell et al., 2019; Hanslmayr et al., 2007; Iemi et al., 2022). Many studies have established a relationship between the amplitude of ongoing activity in the alpha band (α, 8-13 Hz) before stimulus presentation and perceptual performance (Ergenoglu et al., 2004; Hanslmayr et al., 2005; Hanslmayr et al., 2007; van Dijk et al., 2008). Some of these studies have specifically linked response variability in visual tasks to changes in the amplitude of pre-stimulus α-oscillations in the posterior brain regions prior to stimulus presentation (Bompas et al., 2015; Foster et al., 2017; Huang et al., 2019; Kirschfeld, 2008; Lin et al., 2013; Linkenkaer-Hansen et al., 2004; Yang et al., 2014, but see Bays et al., 2015; van Dijk et al., 2008).

Although all of these findings are based on averaging pre-stimulus activity, a few recent studies have provided evidence that oscillatory activity may often consist of burst-like events of high-power neural activity occurring stochastically at different rates, times, and durations (Feingold et al., 2015; Jones, 2016; Lundqvist et al., 2016; Shin et al., 2017; Tinkhauser et al., 2017; van Ede et al., 2018; Zich et al., 2020). An illustrative example is hippocampal theta. Many studies have observed overall increases in average theta power related to navigation in the human hippocampus (Ekstrom et al., 2005; Watrous et al., 2013). However, when looking at single trials in intracranial EEG studies, theta appears in distinct bouts of activity and not (only) as persistent power changes (Goyal et al., 2020). Similarly, α-occipital, one of the most well-known oscillatory characteristics of the human EEG, appears to be expressed in burst-like events (Kosciessa et al., 2020; S. M. Sherman & Guillery, 1998; S. Murray Sherman & Guillery, 2001; van Ede et al., 2018). Nevertheless, α-activity is overwhelmingly studied by averaging many trials, perhaps overlooking important physiological relevant aspects of its temporal, spectral, and temporal structure (Zich et al., 2020). Sustained high power in the averaged spectrum can arise due to increased rates or durations of bursts, power changes of bursts, or an overall increase of power across the spectrum. Thus, traditional trial-averaged analysis can hardly capture this burst-like dynamics. Instead, a trial-by-trial approach can optimally help ascertain burst events (Stokes & Spaak, 2016).

To this end, a real-time approach has the potential to capture a more accurate representation of moment-to-moment variations of the cortical dynamics (Lundqvist & Wutz, 2021) and bring the opportunity to trigger stimuli timed-locked to the occurrence or absence of relevant burst events in the ongoing signal. This approach can provide a robust test of whether burst-like events are associated with particular behavioural outcomes and, in addition, can naturally provide a crucial proof of concept for brain-computer interface (BCI) applications. However, to the best of our knowledge, real-time BCI targeting α-bursts from the human brain has not been addressed so far. The present study reports the outcome of such a novel test using scalp EEG measurements of an ongoing, posterior ⍺-activity based BCI system for real-time burst-triggered stimulus presentation in a visual perception task. So far, α-bursts in the EEG from the occipital cortex have been used as a physiological measure for assessing attentional state and related to errors in driving studies (Papadelis et al., 2007). In addition, EEG α-bursts have been demonstrated to be superior to the EEG α-band power measures in terms of sensitivity and specificity for assessing the driver fatigue under real traffic conditions (Borghini et al., 2014; Simon et al., 2011). Although these kinds of studies mainly use offline data analyses, findings like these have the potential to extrapolate to real-time monitoring/warning systems in real-world scenarios where sustained attention is crucial, such as driving vehicles, piloting aeroplanes, and operating heavy machinery.

Supposing that the accumulation of ⍺-bursts over trials correlates with the behaviour observed in trial-averaged ⍺-power (Lundqvist & Wutz, 2021), then, by specifically targeting the ⍺-burst events, we should see a similar behavioural performance as in trial-averaged studies, but on a much fine-grained scale of oscillatory dynamics of the EEG signal. Based on the empirical evidence and the well-known inhibitory role of ⍺-activity over sensory cortices (Iemi et al., 2022; Jensen & Mazaheri, 2010; Klimesch et al., 2007), we hypothesised that target presentation during the presence of ⍺-bursts would impact subsequent reaction time (RT). In particular, targets presented during ⍺-bursts would lead to slower RT (i.e., worst performance), whereas targets presented outside ⍺-bursts resulted in faster RT (i.e., better performance). This hypothesis, if confirmed, would not only help corroborate and extend ⍺-theories currently based on findings from averaged trials (Bompas et al., 2015; Campagne et al., 2004; Horne & Baulk, 2004; Jung et al., 1997; Kirschfeld, 2008; Klimesch et al., 1998; Lal & Craig, 2002; Lin et al., 2013; Makeig & Inlow, 1993; Makeig & Jung, 1995, 1996; Wyart & Tallon-Baudry, 2009; Yang et al., 2014), but also provide evidence of the putative role of dynamic ⍺-bursts in real-time perception and subsequent behaviour.

We implemented a go/no-go visual detection task in which target presentation was determined in real-time based on the occurrence or the absence of α-bursts in the ongoing occipital EEG signal (**Fig. 1**). We estimated bursts by adapting the eBOSC method (Kosciessa et al., 2020) to the real-time analysis pipeline of the BCI setting and used the last 45 s as the background window, as similarly done in previous applications of the method (Whitten et al., 2011). When a burst event (or its absence) for at least three cyles of the individual alpha frequency (here, Individual Frequency of Interest; IFoI) was detected, the visual stimulus was triggered. In go trials (80%, green light), participants had to respond as fast as possible, whereas in no-go trials (20%, red light), participants had to inhibit the response. Stimuli were randomly and equally delivered during the occurrence/absence of α-bursts, as detected by the BCI (see below for details). Overall, each participant was presented with 240 trials valid for analysis: go/burst (n=96), go/no-burst (n=96), no-go/burst (n=24), and no-go/no-burst (n=24). Note that our task’s stimulus intensity was weaker than regular go/no-go tasks since we wanted to induce a perceptual ambiguity (Benwell et al., 2021) and errors (Chaumon & Busch, 2014). To this end, our study aimed to provide (I) evidence for the functional relevance of oscillatory α-bursts in visual perception and (II) a proof-of-concept for a real-time burst-triggered stimulus presentation BCI setting. The hypothesis, procedure, and analysis pipeline were pre-registered before data collection (https://osf.io/z98ms/). Deviations from the pre-registered procedure are clearly stated in the manuscript. Data and code used in this experiment will be shared upon publication.

**FIGURE 1.**
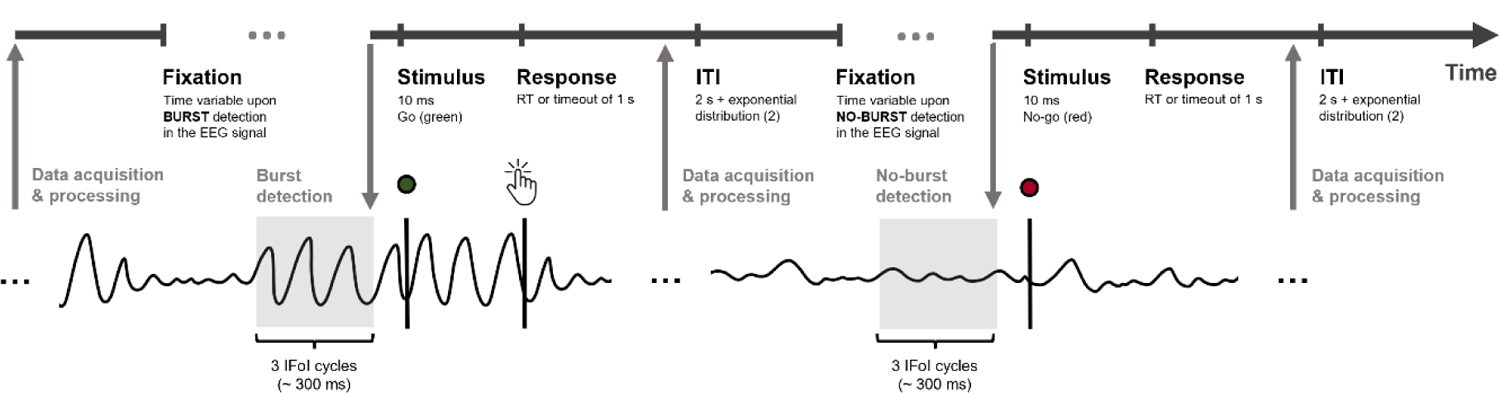
Schema of the dynamics of the sequence of trials in the go/no-go task. Participants had to fixate their gaze on the position of the LED while the BCI system acquired and processed new EEG data. When the pipeline found a (no-)burst – (non-)oscillatory activity for at least three cycles of the individual frequency of interest (IFoI), the stimulus was triggered. In go trials (green light), participants had to respond as fast as they could to the stimuli (with a timeout of 1 s), whereas in no-go trials (red light), participants had to inhibit the response. After the response (or the timeout), the BCI system acquired and processed new data while there was an inter-trial interval (ITI) of 2 s + an exponential distribution (mean = 2 s). Go-stimuli occurred in 80% of trials, with no-go stimuli occurring in 20%. In both types of trials, stimuli were delivered randomly and equally during the occurrence/absence of α-bursts (oscillatory activity) at the target onset. Overall, there are four conditions in our go/no-go task: go/burst (n=96), go/no-burst (n=96), no-go/burst (n=24), and no-go/no-burst (n=24). Here, only go/burst and no-go/no-burst are exemplified.

To anticipate the conclusions, the study showed a connection between oscillatory bursts and behaviour utilising EEG-based BCI system for burst-triggered stimulus presentation in real-time. Stimulus presentation contingent upon the occurrence or absence of occipital α-bursts impacted behaviour (i.e., reaction time and errors) significantly on most participants individually, and as a group.

## RESULTS

### Stimulus presention contingent upon the occurrence or absence of α-bursts impacts subsequent RTs

According to the pre-registered pipeline, the analyses returned a consistent relation between the occurrence/absence of α-bursts in the ongoing occipital EEG signal and the speed of responses to visual targets, both at the individual and the group level. In all 12 participants but two, RTs for go stimuli were slower on average (486 ms; SD = 50 ms) when presented during burst events (as predicted) and faster (486 ms; SD = 50 ms) when presented in the absence of bursts (see **Supplementary Fig. 1** for RT histograms). In five out of those 10 participants, the difference was significant at the individual level (p-values ranged from *<.001* to *.03*; **Supplementary Table 1**). In line with the individual results shown in **Fig. 2**, the group-level difference between burst and no-burst RTs was highly significant (*p = <.001*; **Supplementary Table 1**), reaching a mean difference of 19 ms (SD = 17; Max = 55 ms; Min = −6 ms).

**FIGURE 2.**
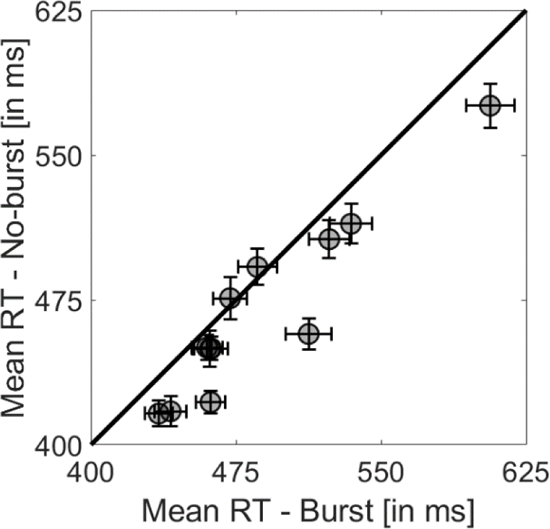
Individual mean reaction times (RT) for burst (x-axis) and no-burst (y-axis) trials for validated trials. Each circle denotes a participant. Horizontal error bars denote the SEM of burst trials, and vertical error bars denote the SEM of no-burst trials. Points below the diagonal line denote that no-burst mean RTs are faster than burst mean RTs. Symbol overlap is coded darker.

### Interim discussion and reality checks

Compared to offline experimental approaches where analysis parameters can be adjusted retrospectively, real-time settings imply a priori parameter choices to constrain the hypothesis and offer more explanatory power. As pre-registered, we performed some reality checks offline to ensure that the a priori choices in the real-time setting conform to expectations. We focused on three critical aspects: first, we checked that we had triggered the stimuli during intended (no-)burst events in the real-time experiment, using the recorded data; second, we checked whether the choice of the IFoI based on training EEG data was representative of the dominant α-frequency during the task; and finally, we checked that the electrode choice (Oz) for the real-time α-burst estimation was representative of α-activity of interest in a larger occipito-parietal (OP) cluster of electrodes.

### Reality check 1: True detection of burst events

We checked, retrospectively, whether real-time target presentation truly occurred during periods of occurrence or absence of α-burst events (see definition in *Methods*). We divided the estimated burst and no-burst trials from the dataset and epoched the data from −45 s to 2 s from stimulus onset (a larger window than the real-time analysis that included the post-stimulus interval). With these, we replicated offline the same analysis performed in real-time (see *Supplementary Materials* for more details). Note that the analysis was applied at the single-trial level in the real-time study, whereas the offline analyses in this section also apply trial-averaged analysis (comparable with the existing traditional literature). For all participants, we calculated the mean power at the time-window of interest (i.e., three-cycles prior to stimulus; TWoI) for burst (Mean = 5.83 µV^2^; SD = 2.75 µV^2^) and no-burst (Mean = 0.18 µV^2^; SD = 0.08 µV^2^) trials (**Fig. 3a**, **Supplementary Fig. 2**, and **Supplementary Table 2**). Overall, the mean power difference between burst and no-burst was 5.65 µV^2^ (SD = 2.69; Max = 12.58 µV^2^; Min = 2.73 µV^2^). We applied a one-tailed t-test (independent samples) with α-level = 0.05 to the mean power values at the TWoI between burst and no-burst trials and found that the two distributions were significantly different from each other (all participants *p = <.001*; **Supplementary Table 2**). Although the statistical analysis was performed within the TWoI, we decided to plot the average of the trials on a wider time window around stimulus onset (−2 to 1s) to visualise the difference in mean α-power between conditions. In **Fig. 3b**, the individual representative plot from one participant (P09) shows a clear difference between the mean (SD) power across conditions (see **Supplementary Fig. 3** for all individual plots). As reflected by the shaded area, burst trials show larger variability (i.e., SD) than no-burst trials in power within the TWoI. In addition, we related the log-transformed mean power at the TWoI with the RT for no-/burst trials at the individual level and found that log-power distributions of both conditions were separated (see **Supplementary Fig. 4** for individual figures). Finally, we checked the amplitude thresholds and demonstrated that they were correctly adjudicated during the study according to the ongoing α-burst activity (see **Supplementary Fig. 5** for individual figures). These results confirm that our approach successfully identified and separated trials with and without α-bursts in ongoing EEG.

**FIGURE 3.**
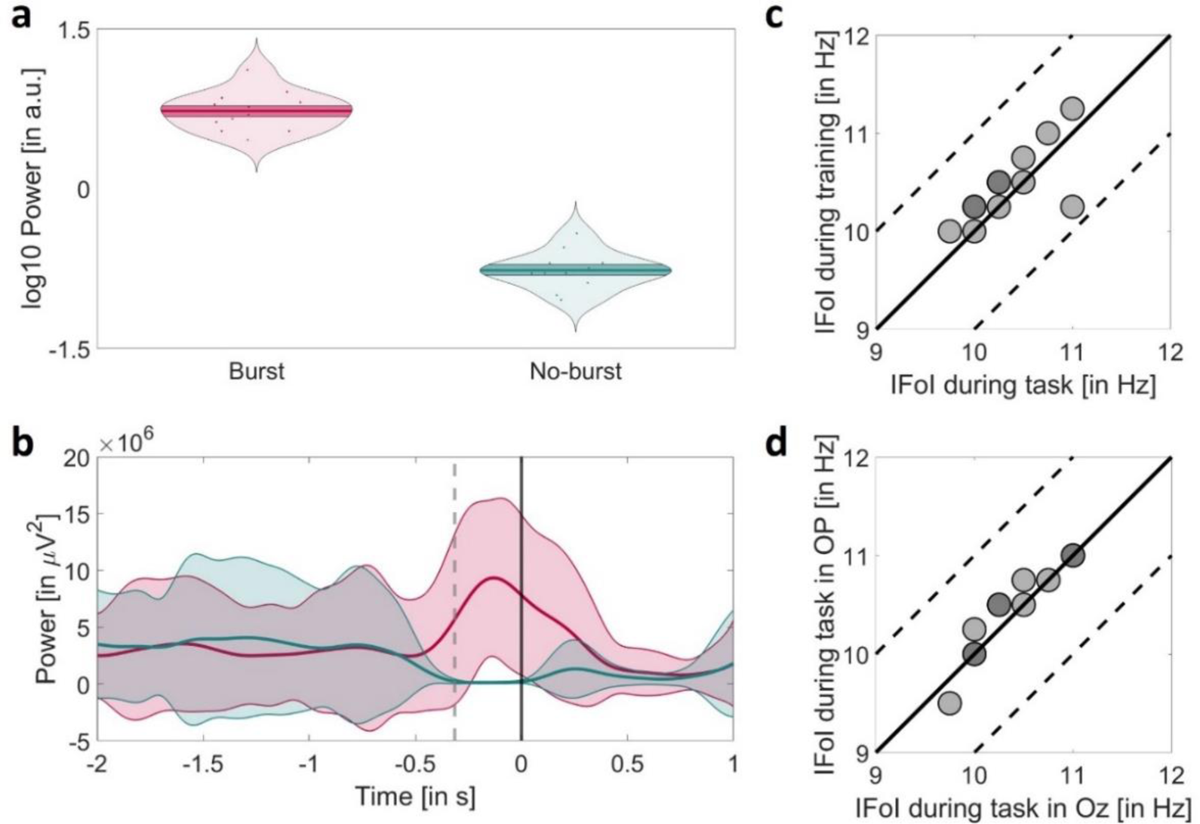
Reality checks of the study. (a) Group-level log-transform mean power at the time window of interest (TWoI) for burst (in red) and no-burst (in green) trials within −2 to 1s from stimulus onset. Solid lines denote the mean power of burst and no-burst trials. Dark shaded areas represent the standard error of the mean interval. Each dot denotes a participant. (b) Individual mean power of burst (in red) and no-burst (in green) trials for participant (P09) within −2 to 1s from stimulus onset. Solid lines denote the mean power of burst and no-burst trials. Shaded areas represent the standard deviation (SD) interval. Solid vertical line denotes the stimulus onset, and dotted vertical lines denote the window of interest (i.e., the last three cycles of IFoI) from stimulus onset. (c) Comparison of individual IFoI during the task [in Hz] and IFoI during the training [in Hz] using Oz electrode. Dashed lines denote ±1 Hz, and each dot denotes a participant. Dot overlap is coded darker. (d) Comparison of individual IFoI during the task [in Hz] using Oz electrode and IFoI during the task [in Hz] using OP-cluster of electrodes. Dashed lines denote ±1 Hz, and each dot denotes a participant. Dot overlap is coded darker.

### Reality check 2: Selection of the frequency of interest (IFoI)

Here, we checked for any potential deviations between IFoI used to estimate bursts in real-time during the task extracted from the training and the actual IFoI during the task execution for each participant. During the training, the mean IFoI peak was 10.60 Hz (SD = 0.41 Hz) with a mean amplitude of 5.12 dB (SD = 3.11 dB), whereas, during task execution, the mean IFoI peak was 10.35 Hz (SD = 0.41 Hz) with an amplitude of 5.72 dB (SD = 2.73 dB) (**Fig. 3c**, see **Supplementary Table 3** for individual results). Overall, the mean peak IFoI difference between task and training was −0.25 Hz (SD = 0.11; Max = −0.5 Hz; Min = 0 Hz), with a mean amplitude IFoI difference of 0.61 dB (SD = 1.63; Max = 3.47 dB; Min = 0.22 dB). The variation in frequency from the IFoI at the single-trial level seems negligible from a time-frequency analysis standpoint. Thus, we can conclude that the BCI setting employed in this study successfully centred the spectral analysis around the desired relevant frequency of interest.

### Reality check 3: Representativity of the electrode of interest

Similarly, we had to decide a priori about the electrode/s of interest used in the BCI setting as for the frequency of interest. We chose the Oz electrode, which was convenient to curtail real-time computational delays. However, one potential concern is that the frequency estimated by that single electrode could be unrepresentative of the α-frequency dominant in a wider cluster of occipito-parietal (OP) electrodes. Therefore, we compared the spectral peak activity in Oz-electrode to that of an OP-cluster of electrodes (P7, P3, Pz, P4, P8, O1, Oz, O2) retrospectively (**Fig. 3d**). The comparison process was analogous to the one described for *Reality check 2*. For the OP-cluster, we computed the IFoI peak (Mean = 10.43 Hz; SD = 0.46 Hz) and IFoI amplitude (Mean = 4.65 dB; SD = 3.49 dB) during the task (see **Supplementary Table 3** for individual results). This yielded that the IFoI of the α-bursts estimated from Oz during the real-time experiment were closely representative of the OP-activity.

## EXPLORATORY ANALYSES

### RT fits using the ex-Gaussian function

Here, we explored how the RT distributions differed between burst and no-burst to learn more about the origin of the behavioural differences seen in the main pre-registered analysis. We characterised the RT-distributions by fitting an ex-Gaussian function, a convolution of a Gaussian and an exponential distribution (Hockley, 1984; Luce, 1986; Ratcliff & Murdock, 1976). Whereas the mean (μ) and standard deviation (σ) for the Gaussian part are thought to reflect, but not exclusively, stimulus- or response-related processes (Schmitz & Wilhelm, 2016), the exponential component (τ, which reflects the skewness of the distribution) is sensitive to central attentional processes, especially those that require inhibitory control (McAuley et al., 2006; Shao et al., 2012; Spieler et al., 1996). We used the *exgauss* toolbox in MATLAB (Bram Zandbelt, 2014) to perform the RT fits. For each participant, we performed a one-tailed permutation test (100,000 randomisations) for the μ parameter (the same statistical tests applied in the main analysis of the study under the same hypothesis, see *Statistical analysis*) and a two-tailed permutation test (100,000 randomisations) for σ and τ parameters (since we did not have any hypothesis about them). **Fig. 4a** illustrates the ex-Gaussian fit for a representative participant (P09), including the RT distribution fit and the psychometric curves for burst and no-burst trials (see **Supplementary Fig. 6** for individual figures). The average μ for burst and no-burst trials was 400 ms (SD = 38 ms) and 393 ms (SD = 41 ms), respectively. The mean RT difference between burst and no-burst was 7 ms (SD = 20 ms; Max = 32 ms; Min = −1 ms). Overall, 7 out of twelve participants showed a difference in RTs in the expected direction (see **Supplementary Table 4** for individual results), with 2 out of these showing a significant p-value (*p < .02*) in the μ parameter. However, we did not observe a significant group-level difference between burst and no-burst (*p = .13*). In terms of σ (burst was 39 ms, SD = 10 ms; no-burst was 37 ms, SD = 14 ms), the average difference was not significant at the group level, and only one participant (P07) showed a significant difference with more variability in burst trials compared to no-burst trials (*p = .01*). Finally, for parameter τ, burst was 86 ms (SD = 17 ms) and no-burst 74 ms (SD = 22 ms), leading to a significant group difference of 12 ms (SD = 17; Max = 41 ms; Min = −1 ms; p = *.01*). Three participants (P01, P03, P07) showed a significant p-value (*p < .05*) for the difference in τ (p-values ranged from *.003* to *.02*), in the same direction as the group. Taking these results together seems that the difference we found in the main analysis between burst and no-burst RTs (*p = < .001*) was best captured by the skewness of the distribution reflected in the exponential component (τ). This finding could be related to the sensitivity to central attentional processes and the requirement of inhibitory control (McAuley et al., 2006; Shao et al., 2012; Spieler et al., 1996) for the go/no-go task. However, given the exploratory nature of this analysis and the fact that overall robust RT differences seemed to diffuse across the various parameters, the interpretation of this analysis should be treated with caution.

**FIG. 4.**
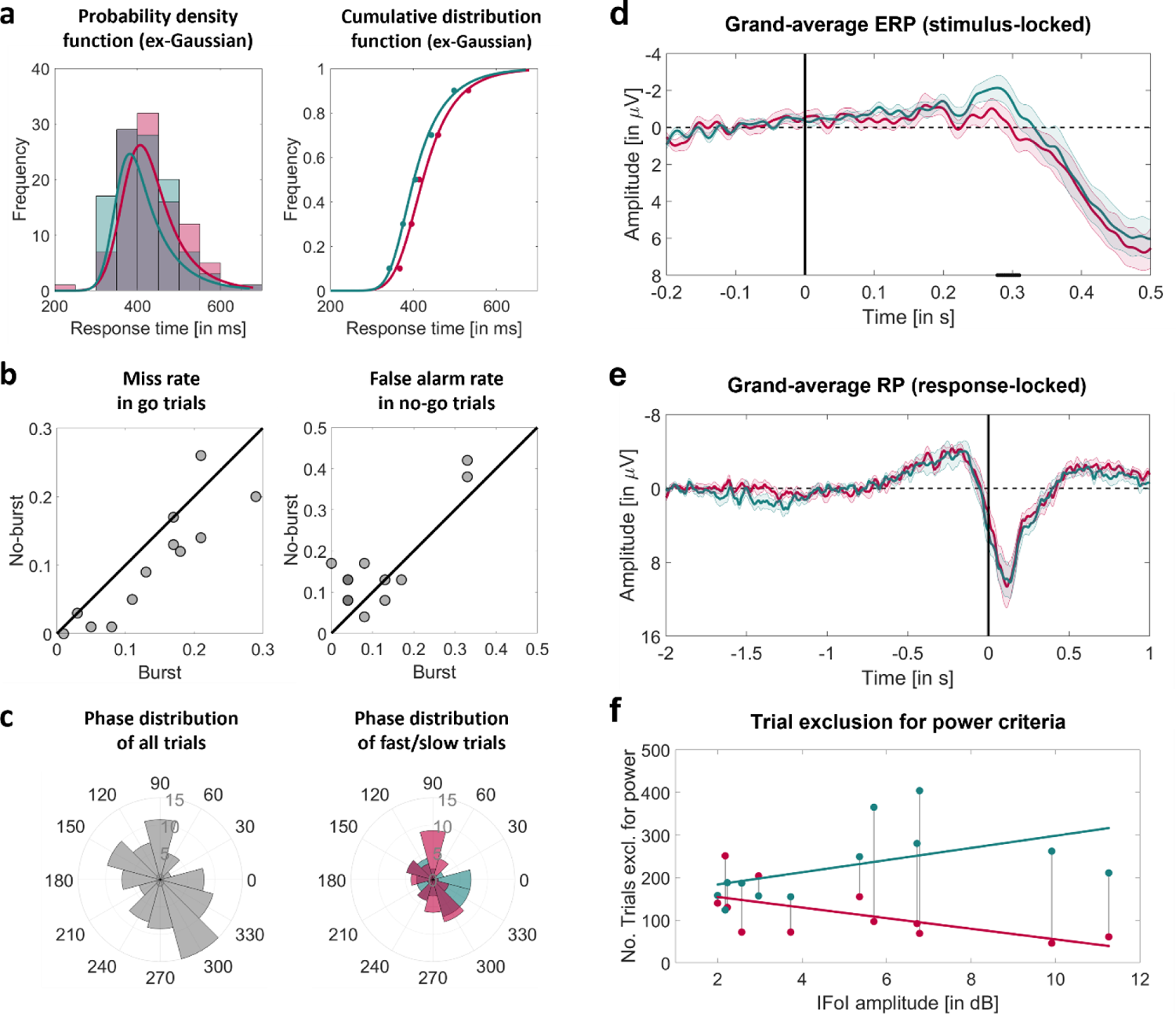
Exploratory analyses of the study. (a) Individual ex-Gaussian fit for burst (in red) and no-burst (in green) trials. For participant P09, the probability density function (PDF) of burst and no-burst trials (on the left) and the cumulative distribution function (CDF) of each trial condition (on the right) are shown. **(b) Comparison of error rates for burst and no-burst trials.** Omission error rates (miss rate) in go trials (on the left) and false alarms or commission error rates in no-go trials (on the right) are shown. Each dot denotes a participant. Points below the diagonal line denote that burst trials contain higher error rates than no-burst trials. Dot overlap is coded darker. **(c) An individual rose plot of the phase distribution at stimulus onset for valid-burst trials (in degrees).** For participant P09, the rose plot of phases for all valid-burst trials (on the left) and the rose plot of fast/slow trials within valid-burst trials (on the right) are shown. Each bin corresponds to 30°**. (d) Grand average ERP of the broad-band activity time-locked to visual stimulus presentation for burst (in red) and no-burst (in green) of go trials from Oz-electrode across time (within −0.2 s to 0.5 s from stimulus onset).** Shaded areas represent the standard error of the mean (SEM) interval. Vertical solid line denotes stimulus onset presentation, and horizontal dashed line denotes zero amplitude. Solid dark line at the bottom of the x-axis denotes the significant cluster of p-values from a paired t-test (α-level = 0.05) for burst and no-burst ERPs over time (p-values were corrected for multiple comparisons via cluster-based permutation test; N = 100,000 randomisations). **(e) Grand average RP of the broad-band activity time-locked to button press for burst (in red) and no-burst (in green) of go trials from Cz-electrode across time (within −1 s to 1 s from response onset).** Shaded areas represent the standard error of the mean (SEM) interval. Vertical solid line denotes response onset, and horizontal dashed line denotes zero amplitude. **(f) Individual IFoI amplitude during the task for burst (in red) and no-burst (in green) trials as a function of the number of trials excluded for power in each condition.** For each participant, the IFoI amplitude during the task (x-axis) presents two related values: number of excluded trials for not satisfying the burst-power condition (dots in red) and the no-burst condition (dots in green). Vertical lines between dots denote the pair of values for each participant. Solid lines denote the linear fit across participants for burst (in red) and no-burst (in green) trials. Group assess of Pearson correlation between IFoI amplitude and number of trials excluded for each participant for burst (left-tailed) and no-burst (right-tail). P-values denote significance in the Pearson correlation at group-level for burst (*ρ = −.66; p = .01*) and no-burst (*ρ = .53; p = .04*) trials.

### Omission and commission errors

Many previous studies optimised to investigate accuracy have found a consistent relationship between high alpha-power and missed targets in detection tasks (Busch et al., 2009; Ergenoglu et al., 2004; Mathewson et al., 2009). Although our study was optimised for RT measurements, we explored the relationship between misses (omission errors in go trials) and burst occurrence. What is more, despite the low number of no-go trials, we reasoned that exploring false alarms (commission errors in no-go trials) during periods with and without burst activity would be informative. If α-bursts indicate periods of lowered sensitivity, one might expect a higher error rate during these episodes (higher α-power) than during no-burst episodes (lower α-power reflecting higher sensitivity). This hypothesis was considered only for go trials (Chaumon & Busch, 2014; Hanslmayr et al., 2007; Klimesch et al., 2007) since we did not have any expectations with no-go trials. In order to increase the number of total trials and capture errors, we decided to use all trials in which stimuli were correctly triggered according to the occurrence of an oscillatory burst event (or its absence) and relax the *RT criterion*. The resulting trials were divided into burst and no-burst conditions for go and no-go trials (**Supplementary Table 5**). For these comparisons, we used a two-tailed paired t-test. The results revealed a significant difference in error rates between burst and no-burst conditions for both go (*t (11) = 3.15, p = .01*) and no-go trials (*t (11) = −2.29, p = .04*) at group-level. In particular, in go trials, omission errors (misses) were more prevalent during a burst episode than during a no-burst (**Fig. 4b, left)**. In no-go trials, commission errors (false alarms) were more prevalent during a no-burst episode than a burst episode (**Fig. 4b, right)**. As expected, bursts (high α-power) were related to higher miss rates in go trials; however, in no-go trials, false alarms were more prevalent during the absence of a burst (low α-power). A key aspect when interpreting these results relies on the differences in cognitive processes between go (i.e., answering to stimuli) and no-go trials (i.e., inhibiting the response to stimuli) and on the assumption that the level of α-power reflects the sensory excitability needed to detect a stimulus (Hanslmayr et al., 2007; Klimesch et al., 2007). In go trials, lower oscillatory α-power (i.e., stronger excitability) may lead to higher detection accuracy and faster reaction times, whereas higher oscillatory α-power (i.e., lower excitability) would impair accuracy (e.g., slower RTs and more false alarms). However, the level of α-power is also thought to reflect the level of inhibition (Klimesch et al., 2007). To this end, in no-go trials, where inhibition is needed to stop the response, higher oscillatory α-power (i.e., stronger inhibition) would help and lead to fewer false alarms. On the contrary, lower oscillatory α-power (i.e., weak inhibition) may lead to answering to visual stimuli producing thereby higher false alarms. We tentatively conclude that our results in the error rate difference are in line with the expected findings and those in the literature, using single-trial and group-averaged pre-stimulus α-power offline.

### Phase-behaviour opposition

Some past and recent studies have addressed the relationship between α-phase and behavioural responses to visual stimuli in humans. Although several studies have related the α-phase of occipito-parietal areas to performance in visual perception (Jensen et al., 2011; Jensen & Mazaheri, 2010; Samaha & Postle, 2015; van Dijk et al., 2008; VanRullen, 2016), other studies have also reported no (O’Hare, 1954; Walsh, 1952) or weakly significant effects (Callaway, 1962; Callaway & Alexander, 1960; Callaway & Yeager, 1960; Dustman & Beck, 1965; Lansing, 1957), and even null evidence for accuracy (Benwell et al., 2017; Ruzzoli et al., 2019) and RTs (Vigué-Guix et al., 2020). Although this analysis was not pre-registered, we explored the potential α-phase/RT relationship at stimulus onset. If the response to a stimulus is related to the α-phase at its onset, we should expect a pattern of opposition within the α-oscillations when comparing slow and fast RT trials at stimulus onset (time = 0; Mathewson et al., 2009). We checked for this opposition pattern using the phase opposition sum (POS) method (VanRullen, 2016). We selected only valid-burst trials (which contain oscillatory activity and a valid response) for this analysis, leading to N = 96 trials for each participant. Trials were median-split into slow (50% RTs) and fast (50% RTs) bins, resulting in N = 48 trials for each condition and participant. EEG data were epoched from −45 to 2 s and then demeaned. No re-referencing was applied. We forward-filtered the signal (electrode Oz) using a Butterworth filter (order 4, one-pass) around the IFoI ± 5 Hz band and computed the phase at stimulus onset using the Hilbert transform. We plotted the distribution phases (**Fig. 4c;** see **Supplementary Fig. 7** and **Supplementary Fig. 8** for the individual phase distribution of all trials and fast/slow trials, respectively) and calculated the averaged phase between fast and slow RTs and their difference in phase using the Circular Statistics Toolbox in MATLAB (Berens, 2009). We applied the POS on phase values for fast/slow trials at individual and group levels. The statistical significance was assessed using nonparametric permutations tests (10,000 iterations/surrogates). Comparing the empirical POS value to surrogate POS distributions dispense with the assumption that the distribution of phases across trials is random and uniform (McLelland et al., 2016). We did not observe any significant group-level effect of POS between slow and fast RTs (*p = .32*) nor at an individual level for any participant (all *ps > .1*; see **Supplementary Table 6** for individual results). The average phase angle difference among participants between slow and fast trials was −43° (SD = 57°). Overall, the results did not return a consistent relationship between the phase of ongoing α at stimulus onset and the speed of responses.

### ERP analyses time-locked to visual stimulus and response

Although the primary aim of this study was to investigate the influence of the ongoing pre-stimulus oscillatory activity on reaction time, we also analysed the event-related potentials (ERPs) to visual stimuli in go trials separately for the burst and no-burst conditions. Previous studies have linked pre-stimulus EEG α-power and the amplitude of visual evoked potentials (Başar et al., 1998; Mazaheri & Jensen, 2008), but have produced mixed effects on different latencies (Başar et al., 1998; Brandt et al., 1991; Ergenoglu et al., 2004; Fellinger et al., 2011; Roberts et al., 2014). Therefore, our assessment here was exploratory. In addition, we did not expect to find identifiable ERP components, given that our stimulus intensity was fairly low by design. We selected only go trials due to the low trial number of no-go events and potential differences in task relevance and motor preparation between the two types of trials. We band-pass filtered the recorded EEG data from electrode Oz using a Butterworth filter (order 4, two-pass, zero-phase) between 0.5 and 40 Hz for each participant. EEG data were re-referenced offline to the left and right mastoids’ average. EEG data were epoched from −200 to 400 ms and then demeaned. All epochs were baseline-corrected with respect to the mean voltage over the 200 ms preceding the onset of stimuli, followed by averaging for burst and no-burst conditions. Since this was an exploratory analysis, we targeted a broad time window, between 200 and 500 ms. For statistical assessment, we performed sample by sample paired t-tests between the mean visual-evoked ERPs at the Oz electrode between burst and no-burst. We used a cluster-based permutation test procedure (100,000 randomisations) to correct p-values (Maris & Oostenveld, 2007; Meyer et al., 2021). **Fig. 4d** shows the (stimulus-locked) ERPs of valid-go trials for burst and no-burst conditions. We found a significant difference between burst and no-burst conditions (*p < .05*) in a 32 ms time window between 280–312 ms after stimulus presentation. Based on previous literature, this period would correspond to the putative N2 (255 – 360 ms) component (Koivisto & Revonsuo, 2003, 2010; Sheldon & Mathewson, 2021). We reckon that this difference in the ERP might reflect the RT-effect found between burst and no-burst trials at group-level in the main analysis. This is in line with the cited previous ERP studies addressing go/no-go tasks, where the larger the peak of the N2 component (here, larger ERP for no-burst), the faster the responses (here, shorter RTs for no-burst events).

Moreover, we decided to analyze the response-locked activity, or readiness potential (RP; Vaughan et al., 1968), for burst and no-burst conditions in go trials. We used the Cz electrode and proproduced the same preprocessing as in the ERP analysis (with t=0 at stimulus onset). EEG data were epoched from −1 to 2 s and then demeaned. All epochs were baseline-corrected using a 200-ms baseline before the stimulus onset. We redefined trials and changed t =0 from stimulus presentation to button response. Trials were epoched from −1 to 1 s (to have the same length), followed by averaging for burst and no-burst conditions. Given that the RP initiates 1-2 s before the movement onset over frontocentral areas reflecting the cortical excitability for the forthcoming movement (Di Russo et al., 2017), we decided to explore the RP differences within a 2s-window previous to response onset. We performed the same statistical assessment as in the ERP analysis. **Fig. 4e** shows the (response-locked) RP of valid-go trials for burst and no-burst conditions. Despite the wave forms display the typical RP on average, no significant differences between burst and no-burst conditions are found (*p > .05*), meaning that perhaps there are no differences in motor areas for the motor preparation to respond to a stimulus between the two conditions. Given that typical RP experiments use protocols rendering much longer RTs, compared to ours, this is not unexpected.

### Exclusion of trials due to power criterion

Although the present study succeeded at detecting ⍺-bursts in the ongoing EEG activity through our custom-built BCI setting (see Reality check 1), the estimation accuracy of (no-)burst varied across participants and conditions, as reflected in the averaged number of trials of 344 (SD = 78; 52%) excluded for the power threshold criterion (see *Supplementary Materials* and **Supplementary Table 7** for more details). For the estimation of burst episodes and the subsequent triggering of the stimuli in real-time, we sought oscillatory activity above a certain power threshold with respect to the overall spectrum for at least three cycles (∼300 ms) of the IFoI of the participant (more details in the *Methods* section). Note that there was an unavoidable time gap between the decision to trigger a stimulus (based on the most updated EEG data) and the actual stimulus presentation due to computational time (∼72 ± 5 ms). For this reason, we sent the stimulus around three-quarters of a cycle ahead of time, and then trials were checked for burst criteria after sending the stimulus for online sorting. We consider this is probably the main reason why an average of 52% of trials were excluded for not satisfying the power criterion at the time of stimulus delivery. We looked for any differences between burst and no-burst trials, and we found that an average of 116 trials were excluded in burst compared to 220 trials excluded in no-burst conditions (see *Supplementary Materials* and **Supplementary Table 8**). At the individual level, 10 out of twelve participants showed more trials excluded in no-burst trials than burst (see **Fig. 4f**). Thus, our criteria made it more difficult to pass a non-burst event during the task execution than to detect a burst of oscillatory α-activity in the ongoing EEG signal. Please note that the criterion for no-burst was not simply the absence of a burst event, but power in the IFoI had to be within the lowest 5%. In addition, we found a significant correlation between the IFoI amplitude during task execution and the number of trials excluded for burst (*ρ = −.62; p = .02*) and no-burst (*ρ = .51; p = <.05*) trials at group-level (see *Supplementary Materials*). During the task, participants with higher IFoI amplitude had fewer trials excluded for bursts than no-bursts. On a general note, we can say that we succeeded in achieving the second main goal of the study: detecting α-bursts in the ongoing EEG activity using a BCI setting for burst-triggered stimulus presentation in a go/no-go task.

## DISCUSSION

This study aimed to harness oscillatory bursts in real-time to predictably modulate behavioural outcomes: the speed of reactions and the likelihood of omission and commission errors. In keeping with this aim, we found that targets presented during ⍺-burst episodes led to slower RTs than those presented outside, leaving an RT difference of 19 ms found between conditions. Regrading errors, targets presented during bursts were more likely to be missed in go trials, whereas in no-go trials, those presented in the absence of bursts led to false alarms more often. Together, we suggest that the behavioural differences found between burst and no-burst conditions appear to unfold over the processing of the target, perhaps at different stages. For example, the fact that the most apparent difference in the ex-Gaussian fit analysis affected the skewness of the distribution, together with the relatively late difference found in the ERP analyses, suggests that at least a part of this behavioural effect would involve late processing stages of decision or response selection. We consider that these effects can relate to the sensitivity of central attentional processes and the requirement of inhibitory control in a go/no-go task, as suggested in previous studies (McAuley et al., 2006; Shao et al., 2012; Spieler et al., 1996). In addition, there is evidence that larger N2 peaks relate to faster RTs (Bahramali et al., 1998; Starr et al., 1995). In the same vein, we observed larger ERP amplitudes in the N2 time window for bursts condition than no-bursts, which relates to our main finding of burst generating faster responses. Overall, we reckon that burst episodes are periods of lowered sensitivity due to the high α-power, which potentially inhibit the cortical response to visual stimuli, with an impact on slower RTs and more misses. On the other hand, no-burst episodes, as periods of higher sensitivity due to low α-power, may facilitate the answering and produce faster RTs at the expense of more false alarms. In addition, when applying the phase opposition sum (POS) method to fast/slow RTs in burst trials, we did not see an effect of α-phase at stimulus onset, suggesting that the RT-effect (found in the main analysis) is principally a result of the oscillatory power of α-bursts.

It is worth mentioning that the first goal of this study was to provide evidence for the intrinsic function of oscillatory α-bursts on reactions to visual events for the possibility to capitalise on this brain-behaviour relationship for an EEG-based BCI application. Prior studies on the ⍺-theories have frequently found a reliable relationship between the pre-stimulus α-power and the behavioural outcome in visual perception using single-trial and within-subject averages (Bompas et al., 2015; Campagne et al., 2004; Horne & Baulk, 2004; Jung et al., 1997; Kirschfeld, 2008; Klimesch et al., 1998; Lal & Craig, 2002; Lin et al., 2013; Makeig & Inlow, 1993; Makeig & Jung, 1995, 1996; Wyart & Tallon-Baudry, 2009; Yang et al., 2014). However, a fair question and a key challenge are to understand whether the behaviour and the sustained oscillatory activity observed in trial-averaged power (Lundqvist & Wutz, 2021) is actually due to the accumulation of transient high-power burst events that happen at different rates, times, and durations from trial-to-trial (Lundqvist & Wutz, 2021; van Ede et al., 2018; Zich et al., 2020). One way to address this challenge is by using a BCI to target the oscillatory dynamics of the EEG signal (reflected in the α-burst events) at the single-trial level. The oscillatory burst event analysis can serve as a sensitive tool to capture single-trial differences, which would go unnoticed with a standard approach of trial-avered power. Such an approach can answer whether a similar neural-behavioural relationship applies to the much more fine-grained scale of moment-to-moment dynamics of the EEG signal and which form it may take (Pernet et al., 2011). As far as we know, we are the first to use this novel approach in targeting, specifically, burst-like events of oscillatory α-activity from the occipito-parietal cortex of human brains in real-time using an EEG-based BCI setting in order to address its link, on a trial-by-trial basis, to reaction times in a go/no-go task. Similar findings are in line with recent studies of oscillatory burst-like events underlying cognitive and motor operations in other frequency bands (Feingold et al., 2015; Lundqvist et al., 2016; M. A. Sherman et al., 2016; Shin et al., 2017; Wutz et al., 2020).

Moreover, it would be important to make explicit in the current formulation of the α-theories whether and how transient events of rhythmic oscillatory activity (bursts) in the ongoing brain activity gate sensory information and shape perception on a trial-by-trial basis. Although the functional inhibition account (Klimesch et al., 2007) is specific to the α-oscillatory activity, it does not distinguish between sustained oscillatory activity and burst events. On that note, Peterson and Voytek (2017) have recently proposed that α-oscillations control cortical gain by modulating the balance between excitatory and inhibitory background activity, and they make the novel prediction that α-activity plays two functional roles: a robust, sustained oscillation mode (>5-10 cycles) that suppresses cortical gain and a weak, bursting mode (of 1-3 cycles) for rapid, temporally-precise gain increases. This view would align with Jensen and Mazaheri (2010) and Mazaheri and Jensen (2010). Together, new models, theories, and analytical methods (Lundqvist & Wutz, 2021) are starting to consider different types of α-activity and assess their respective roles in cognitive processes.

Apart from the theoretical implications, another goal of this study was to provide a proof of concept for using oscillatory α-bursts as a control signal for a BCI. We found that harnessing the detection of neural burst events achieved individual RT differences of up to 55 ms (Mean = 19 ms; Max = 55 ms; Min = −6 ms). One should consider whether such a difference could be meaningful for an EEG-based BCI application from a practical perspective. Note that the potential relevance of such a system does not hinge so much on the average but on single instances. One must consider that time savings can potentially be more considerable on some occasions (for example, looking at the most favourable case, participant P09, one could save as much as 250ms on roughly 10% of the trials). These are substantial results if one considers this is the first proof of concept where many of the parameters and protocol features have been necessarily preset arbitrarily, given the lack of precedent.

One of the major concerns of the study was the high exclusion rate of participants (31 out of 43). However, pre-screening is a common practice in hypothesis-driven real-time BCI studies (Callaway & Yeager, 1960; Lansing, 1957; Vigué-Guix et al., 2020). Note that oscillatory activity in the α-band should be present to establish the role of α-oscillations in subsequent perception. In our study, nearly half of the excluded participants were not either eligible because it was not possible to resolve distinct α-activity from their scalp, or because there was not a unique oscillatory peak within the α-band. The other half of the participants were excluded for not reaching enough α-activity variability to fully account for the two conditions of our study (i.e., low/high oscillatory α-activity). Note that this is also why nearly 50% of trials in the real-time study were rejected because of α-power. Nonetheless, our exclusion rates are comparable with previous studies rejecting almost two-thirds of their participants (Callaway & Yeager, 1960; Vigué-Guix et al., 2020) and even 92 out of 100 participants (Lansing, 1957) for similar reasons. On a general account, it has been thought that individual differences in the measurement of activity with EEG may be partially due to physiological differences (e.g., the thickness of the skull), technical and methodological factors (e.g., type of EEG montage), or specific factors such age, arousal or cognitive demands (Klimesch, 1999).

Although our findings from the study are well in line with a host of older findings relating to α-power and behaviour, the present approach goes beyond those studies in two important ways. First, our study harnessed real-time data analysis trial by trial using an EEG-based BCI system. Second, we focused on bursts of α-activity. Evidence of this kind demonstrates the putative functional role of bursts of oscillatory neural α-activity in occipital areas (Hughes and Crunelli, 2005; Hughes et al., 2011; Lundqvist et al., 2013; Wutz et al., 2020). Moreover, the real-time trial-to-trial nature of this approach provides a potential basis for control signals in EEG-based BCI applications supported by brain-behaviour theories (here, α-theories). For instance, a passive BCI using brain-state dependent stimulation (BSDS) (Jensen et al., 2011) could benefit from our findings and build a BCI application in which stimuli are triggered only in the absence of neural α-burst events. Such BCI would help prevent slower reactions and omission errors. Therefore, this study has shown that it is possible to directly address the connection between oscillatory bursts and behaviour utilising an EEG-based BCI system, allowing for burst-triggered stimulus presentation in real-time. Real-time studies using EEG-based BCI systems are promising research tools than can be used as a test bench for brain-behavioural theories in cognitive neuroscience.

## MATERIALS & METHODS

### Participants

#### Sample size

We set to complete a dataset with N = 12 participants. As per the necessary exclusion requirements set a priori (see below), the attrition rate was high. Of the total 43 participants initially recruited, 16 were discarded for not satisfying the required α-peak criterion in the screening stages (seven did not show a peak at rest, three had double-peak, and six did not show a peak during the task), one was discarded for low discrimination performance, and 14 because of the duration criterion. The final dataset contained EEG and behavioural data from the remaining 12 participants (mean age of 24 years, eight females, all right-handed) without previous history of neurological or psychiatric diseases, with normal or corrected to normal vision, within 18-35 years of age. All participants took part in the study voluntarily after giving informed consent, and they were compensated for their time 10€ per hour. The duration of the experiment varied between 60 and 120 minutes. The study was designed in accordance with the Declaration of Helsinki and was approved by the Institutional Committee for Ethical Review of Projects (CIREP-UPF) (University Pompeu Fabra, Barcelona, Spain) before starting the recruitment. Data from excluded participants were not analysed.

#### Exclusion criteria

A participant was excluded if any of the following criteria were met: *(I) No amplitude peak within the α-band:* This criterion applied to both *screening stages* across the study and ensured that the individual’s endogenous α-oscillation was registered with a sufficiently high signal-to-noise ratio (SNR) to enable the algorithm to classify the (non)occurrence of α-bursts from the ongoing EEG signal. This decision was based on two sub-criteria: strength and uniqueness (see *Screening and estimation of the individual frequency of interest (IFoI)* section for more details). *(II) Time limit of total test duration:* This criterion was applied during the Training block and the Experimental session. Given that the duration of the study depends on the estimation of (non-)occurrence of bursts in the ongoing EEG signal, we decided to establish an objective limit. Thus, we stopped the experiment if a participant spent more than 20 minutes in the Training block or any blocks of the Experimental stage. *(III) False alarms in no-go trials:* This criterion was applied after data collection if a participant had responded in 60% or more trials in the no-go condition. *(IV) Low coefficient of variation (CV) in reaction time (RT):* This criterion was also applied after data collection and ensured that participants had sufficient RT-variability with a minimum CV of 15% (Terentjeviene et al., 2018).

### Experimental procedure and materials

Participants sat on a comfortable chair wearing an EEG cap in front of a green-red bi-colour LED (Manufacturer: Kingbright, Reference: L-59SURKCGKW) at a distance of 90 cm at eye level in a dark and acoustically and electrically attenuated chamber. The LED was attached to a parallel port (forward voltage = 3.84 V, forward current = 0.5 µA) and mounted in a serial circuit with different value resistances depending on the LED colour (one green light resistance = 217 kΩ, and two possible red-light resistances = 933 kΩ or 820 kΩ). The resistance of the green light was permanently fixed, whereas the resistance of the red light was adjusted for each participant to reach the most similar subjective brightness between colours. Note that using different red lights across participants for the no-go trials did not influence the RTs (see *Supplementary Materials* for more details).

During the go/no-go task, visual stimuli were presented to participants via the illumination of the LED either in green or red (10 ms duration), at a time decided from the real-time analysis of the EEG-based BCI setting (see below). Participants were asked to respond with their right finger via a button press as fast as possible each time the LED was lit up in green (go condition) or withhold the response if the LED turned red (no-go condition). Once the response had been given or had reached the timeout with no response (of 1 second), an Inter-Trial-Interval of 2 seconds + exponential distribution (mean of 2 seconds) of time up to 20 seconds started to prevent fixed temporal expectation. Then, a new trial started with the search for a new (no-)burst event. RTs were measured from the onset of the visual target until a button press was detected. Trials were randomised across conditions in both the training and experimental blocks of the study.

The experimental protocol followed four phases. In the *(i) IFoI-rest screening test,* a 5-minutes of resting EEG with closed eyes was recorded to determine the individual frequency of interest (i.e., IFoI-rest) within the α-band (see *Screening and estimation of the individual frequency of interest (IFoI)* for more details). This value was further used in the *(ii) Training block* of 40 trials (identical to the *Experimental blocks,* see below) introduced to familiarise participants with the task and to acquire new data to estimate the IFoI during the task. This procedure considered the potential changes in the individual α-peak (both in amplitude and frequency) between eyes-closed vs eyes-open and between resting-state and in-task mode (Benwell et al., 2019; Samaha & Postle, 2015). The updated value was used in the *(iii) Experimental blocks*, where visual targets were triggered depending on the (non-)occurrence of bursts in the α-activity from the ongoing EEG signal (see *Real-time stimulus presentation*). The experimental session consisted of 6 blocks, each block ending after acquiring a total of 40 valid trials (lasting 14 minutes on average). Each participant completed a total of 240 valid trials, 192 for go-trials (80%) and 48 for no-go trials (20%). Out of these trials, half of the trials were triggered during a burst and half during a no-burst event. The primary measures were reaction times (RT), commission errors (i.e., false alarm responses to no-go stimuli), and omission errors (i.e., misses to go stimuli). After the collection of data for each participant, the *(iv) Post-hoc behavioural screening* was applied for *False alarms in no-go trials* and *Low coefficient of variation (CV) in reaction time (RT)* (see details in *Participants* section).

#### EEG recordings

Continuous EEG data was recorded from 16 passive electrodes (F3, Fz, F4, FC1, FC2, C3, Cz, C4, P7, P3, Pz, P4, P8, O1, Oz, O2) placed according to the 10-20 international system. Additional external electrodes were used for recording horizontal ocular movements (one electrode) and left and right mastoids (two electrodes), placed for offline re-referencing. The AFz electrode was used as the online reference and the right mastoid as the ground electrode. The data was recorded using an ENOBIO 20 5G system at a sampling rate of 500 Hz and a touch-proof medical adapter (all manufactured by *Neuroelectrics, Barcelona, Spain*).

#### Screenings and estimation of the individual frequency of interest (IFoI)

The EEG data was filtered by applying a Notch filter at 50 Hz, a high-pass filter using a second-order Butterworth at 0.5 Hz, and a low-pass filter using an eighth order Butterworth filter, and data was linearly demeaned. We estimated the power spectrum density within the α-band (5-15 Hz) from the Oz-electrode using the Welch method (window = 500 ms; overlap = 10%; resolution = 0.25 Hz). Power spectrum was averaged across the electrodes of interest for each participant and normalised by the mean power spectrum from 0.5 to 45 Hz. We verified that the strength of the peak (power at the local maximum within the 5 to 15 Hz window) was greater than average power in the 0.5 to 45 Hz window. If a single frequency peak existed within a ±5 Hz band from the IFoI peak, it was considered the *individual frequency of interest (IFoI)* and used later as a parameter for the real-time analyses. Otherwise, participants were excluded (see *Exclusion Criterion*).

#### Real-time triggering of visual stimuli during α-bursts

We developed a real-time EEG-based BCI setting using custom-written code in MATLAB *(The MathWorks Inc., Natick, MA, USA)* and the Lab Streaming Layer (LSL) library *(Swartz Center for Computational Neuroscience, UCSD, USA)*. We adapted the BCI setting from Vigué-Guix et al. (2020) and designed a new setting to trigger visual stimuli at the (non-)occurrence of α-bursts in real-time estimated from electrode Oz. We also built a GUI in MATLAB to keep track of the experiment on a trial-by-trial basis. To trigger a visual target, the BCI setting iterated through the following steps (see *Supplementary Materials* for the detailed algorithm): (I) *Data acquisition* of a 45-second sliding window of the most up-to-date data (Whitten et al., 2011); (II) *Data reflection* of the beginning and end of the window; (III) *Data filtering and demeaning* with a band-pass forward filter of 4th-order Butterworth between 0.5 and 45-Hz; (IV) *Time-frequency analysis* using 6-cycle Morlet wavelets within 2 to 38 Hz; (V) *Data trimming* with reflected edges were removed; (VI) *Log(frequency)-log(power) fitting* of the wavelet-derived power spectrum using a robust regression; (VII) *Threshold estimations* of *artifact, burst power, no-burst power,* and *duration threshold*; (VIII) *Checking necessary conditions for triggering stimulus* in terms of power and duration criteria (if conditions were not met, loop went back to step (I)); (IX) *Stimulus presentation* of visual targets (go or no-go, depending on condition); (X) *Response* collection or timeout of 1-second; (XI) *Data acquisition update* of the 45-second window with the most up-to-date data, and repetition of the (I-V) steps. (XII) *RT criterion* check (RT within 50 and 1000 ms, only applied in the go condition), *burst criterion check* (90% of data points of the last three cycles had to be higher than the *burst power threshold* and lower than the *artefact power threshold*), or *No-burst criterion check* (90% of data points of the last three cycles had to be lower than the *no-burst power threshold* and lower than the *artefact threshold*); (XIII) *Trial counter* of the number of valid trials and continued with the next iteration until reaching the number of trials of a block (N = 40). Note that if a step/criterion was not satisfied at any point of the iteration, the BCI setting started a new iteration from step (I). Participants had a break between blocks, and a new block began with the BCI setting starting from step (I).

### Statistical analyses

Only valid go trials of each α-burst condition were included in the analyses and for all tests α-level was 0.05, unless otherwise indicated. Given that reaction time data do not follow a normal distribution but a positive right-skewed distribution, the statistical significance was assessed using nonparametric permutations tests (Ernst, 2004; Morís Fernández & Vadillo, 2020) both at individual and group levels. For the *individual (single-participant) statistical tests*, we tested the difference of the mean-RT distributions between burst and no-burst using a Monte Carlo randomisation procedure (100,000 randomisations) individually for each participant (i.e., one-tailed permutation test) to obtain a p-value associated with the observed mean-RT difference. In the *group statistical tests*, we performed the same procedure and estimated the p-value of the inter-individual mean burst/no-burst difference of RTs for all participants.

## Acknowledgements

This research was supported by the *Ministerio de Economia y Competitividad* ((PSI2016-75558-P and PID2019-108531GB-I00, both to S.S.F.). We thank Manuela Ruzzoli and Lena Matyjek for the insightful comments on the manuscript.

## Author Contributions

IVG: Conceptualisation Methodology, Software, Validation, Formal analysis, Investigation, Data curation, Writing Original Draft & Editing, Visualisation. SSF: Conceptualisation, Writing Original Draft & Editing, Supervision, Funding acquisition.

## Data Accessibility Statement

Data will be uploaded in Open Science Foundation (OSF) and made public upon publication.

## Abbreviations

BCI: Brain-Computer Interface

RT: Reaction time

MEG: Magnetoencephalography

EEG: Electroencephalography

OSF: Open Science Foundation

ITI: Inter-Trial Interval

OP: Occipito-parietal

IFoI: Individual Frequency of Interest

LSL: Lab Streaming Layer

POS: Phase Opposition Sum

BSDS: Brain-State Dependent Stimulation

