## Supplementary Materials for "Using occipital ⍺-bursts to modulate behaviour in real-time"

#### METHODS: Algorithm for real-time burst-triggering of stimuli

To trigger a visual flash (target), the BCI setting iterated through the following steps:

1. *Data acquisition.* A 45-second sliding window (containing 22500 data points) was used to update an EEG data buffer with the most recent available data from the Oz-electrode and used for the estimate of background power (as in Whitten et al., 2011).
2. *Data reflection.* The first and last 2 s of data from the 45-s window were reversed in time and concatenated to the beginning and end (respectively) of the data buffer, leading to a 49-second window. This procedure attenuated edge artifacts from filtering and computing time-frequency analysis (Cohen, 2014).
3. *Data filtering.* Data in the 49-second window was band-pass forward filtered between 0.5 and 45-Hz with a 4th-order Butterworth filter, and data was demeaned using MATLAB built-in functions.
4. *Time-frequency analysis.* Time-frequency transformation of the 49-second window was performed using 6-cycle Morlet wavelets (Grossmann & Morlet, 1984) with 18 logarithmically spaced centre frequencies (as in Whitten et al., 2011), the IFoI was the 10th frequency and the approximate frequency-band ranged from 2 to 38 Hz.
5. *Data trimming.* EEG data and wavelet-derived power from the 49-second window was trimmed and reflected edges were removed, leading to the original 45-second window of the data buffer (Cohen, 2014).
6. *Log(frequency)-log(power) fitting.* The wavelet-derived power spectrum was linearly fit in log(frequency)-log(power) coordinates using a robust regression with bisquare weighting (Holland & Welsch, 1977; Kosciessa et al., 2020) to improve the linear fit of the background spectrum (Donoghue et al., 2020), with the underlying assumption that the EEG background spectrum is characterized by coloured noise of the form $A*f(-\alpha)$ (Buzsáki & Mizuseki, 2014; He et al., 2010; Linkenkaer-Hansen et al., 2001). To further improve the linear fit of the background spectrum with the robust regression (Donoghue et al., 2020), power estimates within a wavelet passband around the IFoI (i.e., IFoI ± 1) were removed prior to fitting.
7. *Threshold estimations.* Power thresholds for rhythmicity (i.e., the occurrence of α-bursts) and non-rhythmicity (i.e., non-oscillatory signal) at each frequency was set at different percentiles of $\chi^2(2)$-distribution of power values, centred on the linearly fitted

estimate of background power at the relevant frequency (for details see Whitten et al., 2011):

- *Artifact threshold:* The power value was set at the 95th percentile of a $\chi^2(2)$-distribution of power values centred at the slowest frequency (around ~2 Hz) to discern between the true α-bursts and high-amplitude in the EEG signal due to eye blinks and muscle artifacts.
- *Burst power threshold:* The power value was set at the 95th percentile of a $\chi^2(2)$-distribution of power values centred at the IFoI-task frequency.
- *No-burst power threshold:* The power value was set at the 50th percentile of a $\chi^2(2)$-distribution of power values centred at the IFoI-task frequency.
- *Duration threshold:* A theoretical duration threshold of a minimum duration of 2.5 cycles (i.e., ~200 ms prestimulus threshold-window containing 100 data points) of the IFoI was used and set at the end of the 49-second window.

8. *Checking necessary conditions for triggering stimulus.* Both power and duration criteria had to satisfy the necessary conditions for triggering stimulus depending on the trial EEG activity-type (i.e., burst/no-burst):
    - *Burst trials:* All data points from the prestimulus threshold-window had to be higher than *Burst power threshold* and lower than *Artifact threshold*.
    - *No-burst trials:* All data points from the prestimulus threshold-window had to be lower than *No-burst power threshold* and lower than *Artifact threshold*.

   If in a given trial these criteria were met, the BCI setting continued and triggered the corresponding visual stimuli for the trial-type; otherwise, the BCI setting algorithm returned to step (1) and updated the initial EEG data buffer.
9. *Stimulus presentation.* Visual targets (go or no-go, depending on condition) were delivered after the validation depending on the trial-type.
10. *Button response*. Reaction time was collected from a button press via a response box connected to the parallel port. If no response was given by the participant (because of a no-go trial or a missed go trial), then the time-out of 1-second was reached and the BCI setting iterated towards the next step.
11. *Data acquisition update.* After the response, we updated the 45-second window with the most recent available data, as in step (1), and re-did the following steps until reaching again step (5).
12. *Trial validity criterion*. Check of trial validity with the updated data. This criterion was set in order to exclude trials in real-time and ensure that (no-)burst activity conditions were met during the computational time of the pipeline (~72 ± 5 ms) between acquisition of data in step (1) and stimulus presentation in step (9). Only trials that satisfied the following criteria were accepted as valid:

- *(No-)burst criterion:*
    - *True detection of burst event:* 90% of data points of the last 3 cycles (~300 ms) of the IFoI before stimulus onset had to be higher than the *Burst power threshold* and lower than the *Artifact threshold*, both thresholds determined at step (7).
    - *True-detection of no-burst event:* 90% of data points of the last 3 cycles (~300 ms) of the IFoI before stimulus onset had to be lower than the *No-burst power threshold* and lower than the *Artifact threshold*, both thresholds determined at step (7).
- *Reaction time criterion*: RT within 50 and 1000 ms (only applied in the go condition)[1].

13. *Trial counter.* If the intended number of trials was reached in a given block (N=40), the algorithm of the BCI setting stopped; otherwise, another iteration started until the desired number of trials per condition was collected. In between blocks, participants had a break and a new block begun with the BCI setting starting from step (1).

Note that if at any point of the iteration a step/criterion was not satisfied, the BCI setting started a new iteration from step (1).

[1] Since the *Reaction time criterion* only applied to go trials, some no-go trials with actual responses were accepted as valid trials in this section (*see Exploratory Analysis* for further details).

### METHODS: Different red lights in no-go trials across participants

Given that participants had two types of red-lights to adjust the intensity similarly to the green-light, we wanted to assess whether there was any relationship between the red-light used for the participants and their average reaction time (RT) in the task (**Supplementary Table 2**). We computed a t-test (independent samples) with α-level = 0.05 to compare the mean RTs between the first six participants (P01-P06; red light with 933 kΩ) and from the last six participants (P07-P012; red light with 820 kΩ). The result was not significant (*t (10) = 0.34, p = .74, dz = .20*). Thus, we can conclude that using different red lights across participants for the no-go trials did not influence the RTs.

### RESULTS: Individual RTs for burst and no-burst trials

**SUPPLEMENTARY TABLE 1. Individual data of reaction time (RT) for burst and no-burst trials.** For each participant, the mean (SD) RT of all trials, the Coefficient of Variation (CV), the mean (SD) RT for burst trials, the mean (SD) RT for no-burst trials, and the difference in mean RT between burst and no-burst. Individuals assess of statistical difference in the RT from the one-tailed permutation test (p<0.05). P-values in bold denote the significance difference in the RT between burst and no-burst trials for 5 participants (P07, P09, P10, P11, P12). RT difference in bold highlight the results that go in line with our hypothesis.

| Part. | Mean (SD) RT (all trials) [in ms] | CV RT | Mean (SD) RT burst [in ms] | Mean (SD) RT no-burst [in ms] | RT diff. [in ms] | Statistics | |
|---|---|---|---|---|---|---|---|
| | | | | | | p | d |
| **1** | 457 (74) | 0.16 | 462 (84) | 450 (59) | **12** | .12 | 0.17 |
| **2** | 525 (104) | 0.20 | 534 (107) | 514 (101) | **20** | .09 | 0.19 |
| **3** | 456 (73) | 0.16 | 460 (80) | 450 (64) | **10** | .18 | 0.13 |
| **4** | 514 (99) | 0.19 | 523 (104) | 506 (95) | **17** | .12 | 0.17 |
| **5** | 490 (97) | 0.20 | 486 (99) | 492 (91) | -6 | .67 | -0.06 |
| **6** | 458 (92) | 0.20 | 461 (90) | 450 (91) | **12** | .18 | 0.13 |
| **7** | 592 (118) | 0.20 | 606 (123) | 576 (111) | **31** | **.03** | 0.26 |
| **8** | 453 (99) | 0.22 | 451 (95) | 454 (103) | -2 | .61 | -0.04 |
| **9** | 428 (69) | 0.16 | 435 (70) | 416 (66) | **19** | **.03** | 0.28 |
| **10** | 486 (101) | 0.21 | 513 (115) | 457 (79) | **55** | **<.001** | 0.56 |
| **11** | 450 (81) | 0.18 | 462 (76) | 422 (56) | **40** | **<.001** | 0.60 |
| **12** | 433 (84) | 0.19 | 441 (82) | 417 (75) | **24** | **.02** | 0.30 |
| **Mean (SD)** | **479 (91)** | **0.19 (0.02)** | **486 (94)** | **467 (83)** | **19 (17)** | **<.001** | **0.22 (0.20)** |

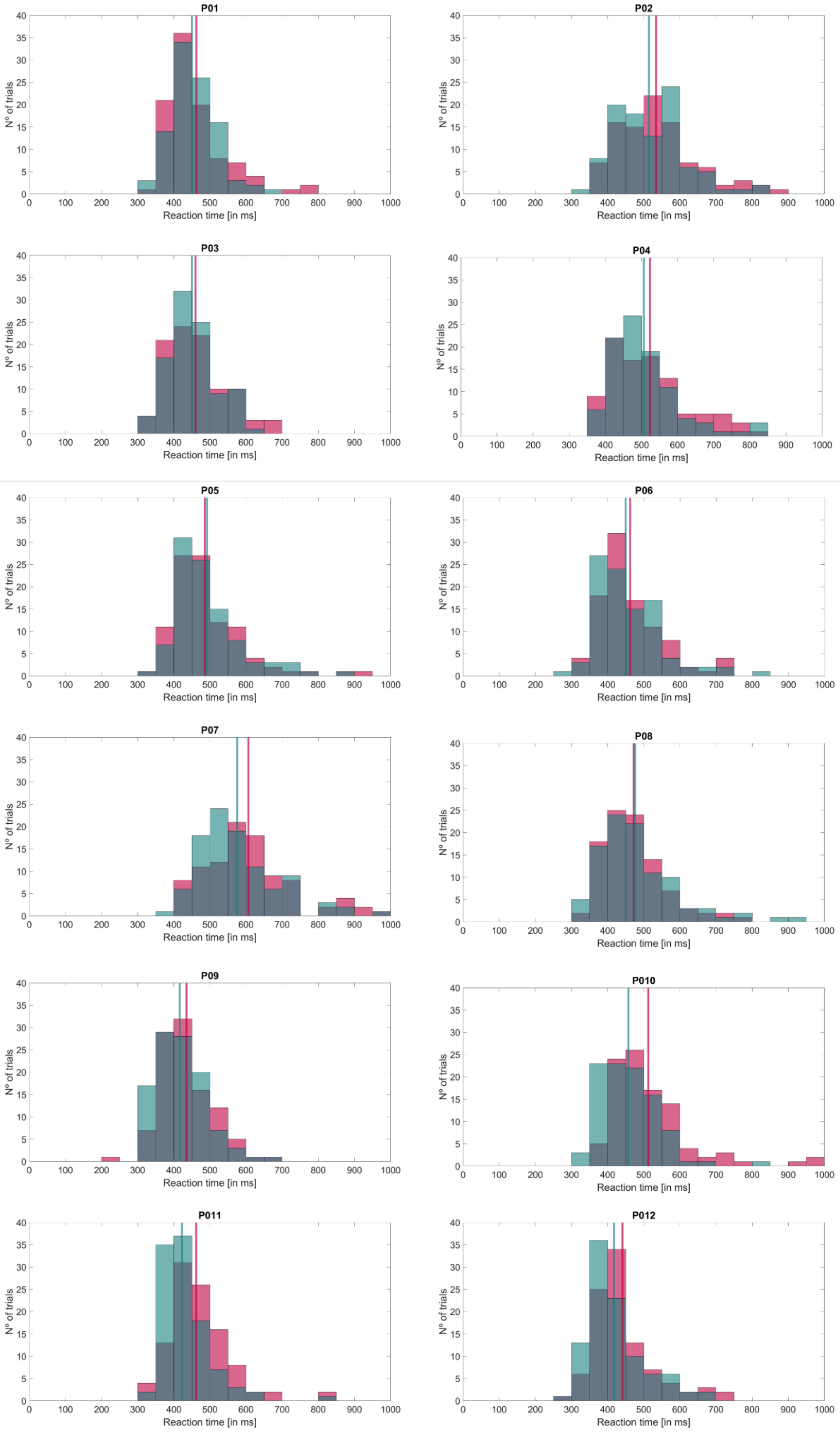


**SUPPLEMENTARY FIGURE 1. Individual histogram of reaction times (RT) for burst (in red) and no-burst (in green) for all validated trials [in ms].** Vertical solid lines denote the mean RT of burst (in red) and no-burst trials (in green).

### REALITY CHECK: True detection of (no-)bursts

**SUPPLEMENTARY TABLE 2. Individual power at the time-window of interest (TWoI) and amplitude thresholds data for burst and no-burst trials.** For each participant, the mean (SD) pre-stimulus α-power [in μV$^2$] for burst and no-burst trials. Individual t-tests assess statistical difference in the α-power and amplitude thresholds ($p<.05$) and the effect size (Cohen's d; d) is also provided. All participants showed a significant difference ($p<.001$) in both the α-power and the amplitude thresholds between burst and no-burst.

| | **POWER TWoI** | | | | | **AMPLITUDE THRESHOLDS** | | | | |
|---|---|---|---|---|---|---|---|---|---|---|
| **Part.** | **Mean (SD) burst [in μV$^2$]** | **Mean (SD) no-burst [in μV$^2$]** | **Diff. power [in μV$^2$]** | **d** | **p** | **Mean (SD) burst [in μV]** | **Mean (SD) no-burst [in μV]** | **Diff. amp. Th. [in μV]** | **d** | **p** |
| 1 | 3.46 (1.77) | 0.16 (0.09) | 3.30 | 2.63 | **<.001** | 1.23 (0.23) | 0.28 (0.05) | 0.94 | 5.66 | **<.001** |
| 2 | 6.14 (2.71) | 0.28 (0.15) | 5.86 | 3.05 | **<.001** | 2.75 (0.89) | 0.64 (0.21) | 2.11 | 3.27 | **<.001** |
| 3 | 7.08 (5.02) | 0.13 (0.07) | 6.96 | 1.96 | **<.001** | 1.26 (0.42) | 0.29 (0.10) | 0.97 | 3.19 | **<.001** |
| 4 | 4.93 (2.93) | 0.09 (0.04) | 4.84 | 2.34 | **<.001** | 0.80 (0.15) | 0.20 (0.04) | 0.68 | 6.02 | **<.001** |
| 5 | 4.52 (1.91) | 0.18 (0.11) | 4.34 | 3.21 | **<.001** | 1.70 (0.25) | 0.39 (0.06) | 1.31 | 7.13 | **<.001** |
| 6 | 6.45 (3.71) | 0.20 (0.13) | 6.25 | 2.38 | **<.001** | 2.13 (0.62) | 0.49 (0.14) | 1.64 | 3.66 | **<.001** |
| 7 | 2.83 (1.33) | 0.10 (0.07) | 2.73 | 2.91 | **<.001** | 1.08 (0.36) | 0.25 (0.08) | 0.83 | 3.20 | **<.001** |
| 8 | 4.21 (2.39) | 0.16 (0.09) | 4.05 | 2.40 | **<.001** | 1.36 (0.27) | 0.32 (0.06) | 1.05 | 5.29 | **<.001** |
| 9 | 8.08 (5.55) | 0.20 (0.14) | 7.89 | 2.01 | **<.001** | 1.78 (0.22) | 0.41 (0.05) | 1.37 | 8.45 | **<.001** |
| 10 | 3.45 (1.27) | 0.17 (0.08) | 3.27 | 3.63 | **<.001** | 1.86 (0.46) | 0.43 (0.11) | 1.43 | 4.23 | **<.001** |
| 11 | 5.84 (2.65) | 0.16 (0.10) | 5.68 | 3.03 | **<.001** | 1.57 (0.37) | 0.36 (0.09) | 1.21 | 4.46 | **<.001** |
| 12 | 12.96 (7.17) | 0.38 (0.24) | 12.58 | 2.48 | **<.001** | 3.13 (0.84) | 0.72 (0.19) | 2.41 | 3.97 | **<.001** |
| **Mean (SD)** | **5.83 (2.75)** | **0.18 (0.08)** | **5.65 (2.69)** | **2.10** | **-** | **1.72 (0.68)** | **0.40 (0.16)** | **1.33 (0.52)** | **2.52** | **-** |

**Supplementary Fig. 3** shows the violin plots of the overall mean of power within the time-window of interest (TWoI) for burst and no-burst trials individually for each participant. We also plotted the average of the trials around stimulus onset (-2 to 1s) to visualize the difference in mean power between burst and no-burst conditions. In **Supplementary Fig. 4,** all the individual plots show a clearly a visual difference between the mean (SD) power across conditions. In particular, burst trials show a larger variability (i.e., SD) in power reflected in the shaded area compared to no-burst trials within the TWoI.

Furthermore, we decided to see the relationship between the log-transformed mean power of the TWoI and the reaction time (RT) for burst (in red) and no-burst (in green) trials at individual level. **Supplementary Fig. 5** shows that there is a clear distinction between our independent variable (i.e., the log-transformed mean power) for burst and no-burst trials, whereas the difference between the dependent variable (i.e., RTs) varies across participants.

Finally, we checked that amplitude thresholds were correctly adjudicated according to the ongoing α-burst activity. We calculated the amplitude thresholds across participants for burst (Mean = 1.72 μV; SD = 0.68 μV) and no-burst (Mean = 0.40 μV; SD = 0.16 μV) trials and also for each participant (see **Supplementary Table 4**). Overall, the mean amplitude difference between burst and no-burst thresholds was 1.33 μV (SD = 0.52; Max = 2.41 μV; Min = 0.68 μV). We assessed the difference between burst and no-burst amplitude thresholds by applying a one-tailed t-test (independent samples) with α-level = .05. All participants showed a significant p-value (*p = <.001*; **Supplementary Table 4**, **Supplementary Fig. 6**).

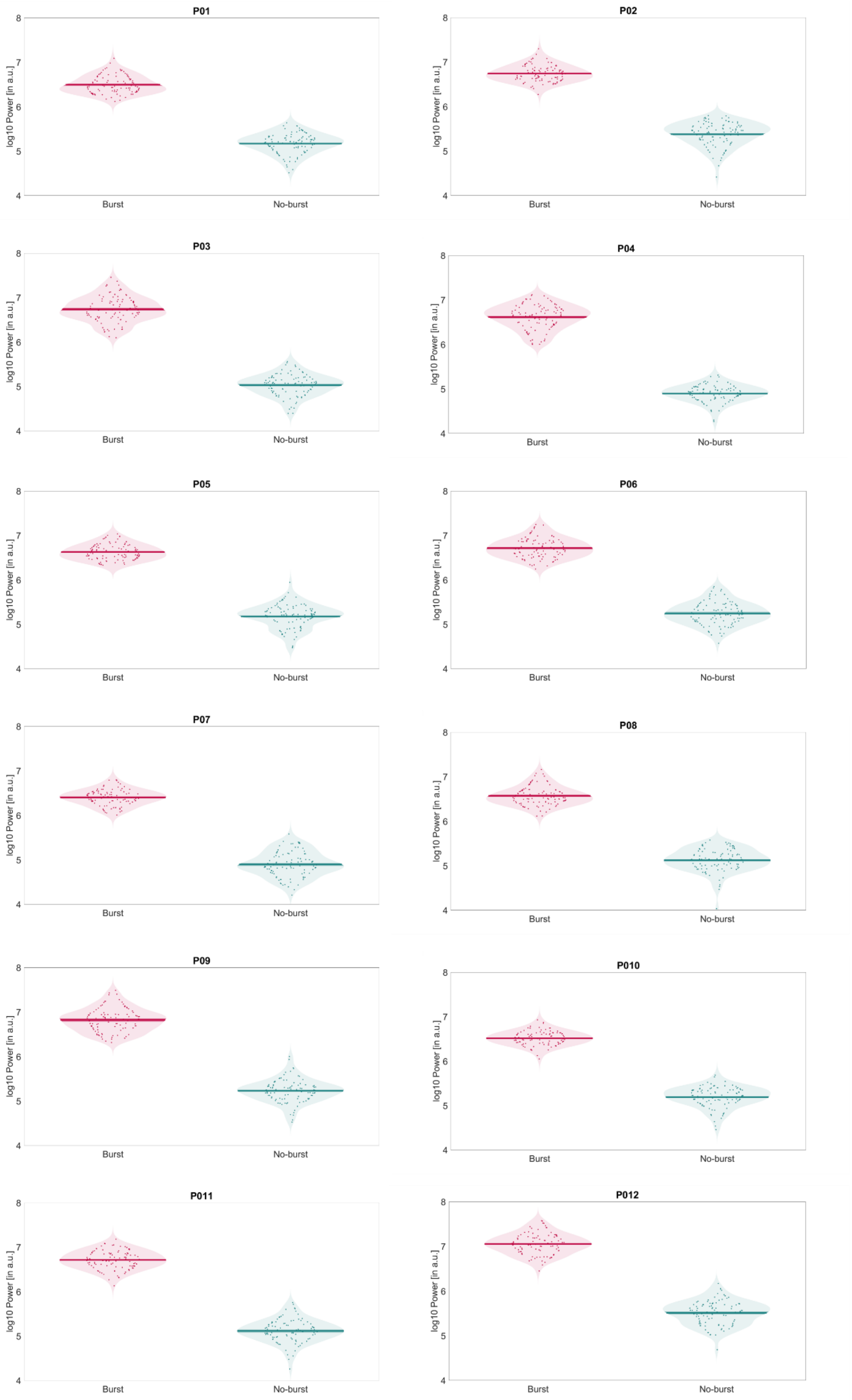


**SUPPLEMENTARY FIGURE 2. Individual violin plots of the mean power for burst (in red) and no-burst (in green) for validated trials.** Violin plots include the mean ± SEM power at individual level. Dots denote the mean power at the time-window of interest (TWoI, i.e., the last 3 cycles of IFoI) for each trial.

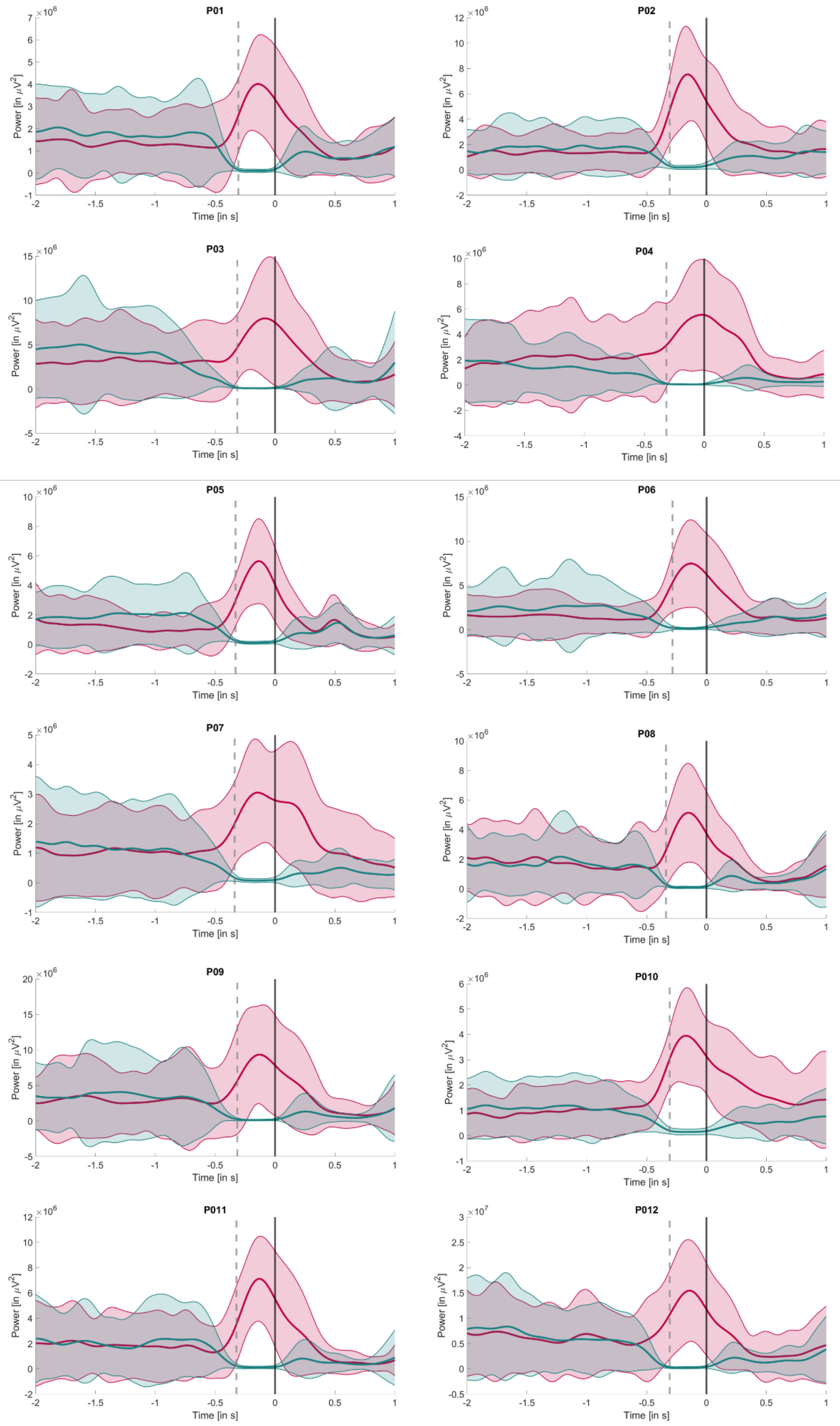


**SUPPLEMENTARY FIGURE 3. Individual mean power of burst (in red) and no-burst (in green) trials at individual level within -2 to 1s from stimulus onset.** Solid lines denote the mean power of burst and no-burst trials. Shaded areas represent the standard error of the mean (SEM) interval. Solid vertical line denotes the stimulus onset and dotted vertical lines denotes the time-window of interest (TWoI) from stimulus onset.

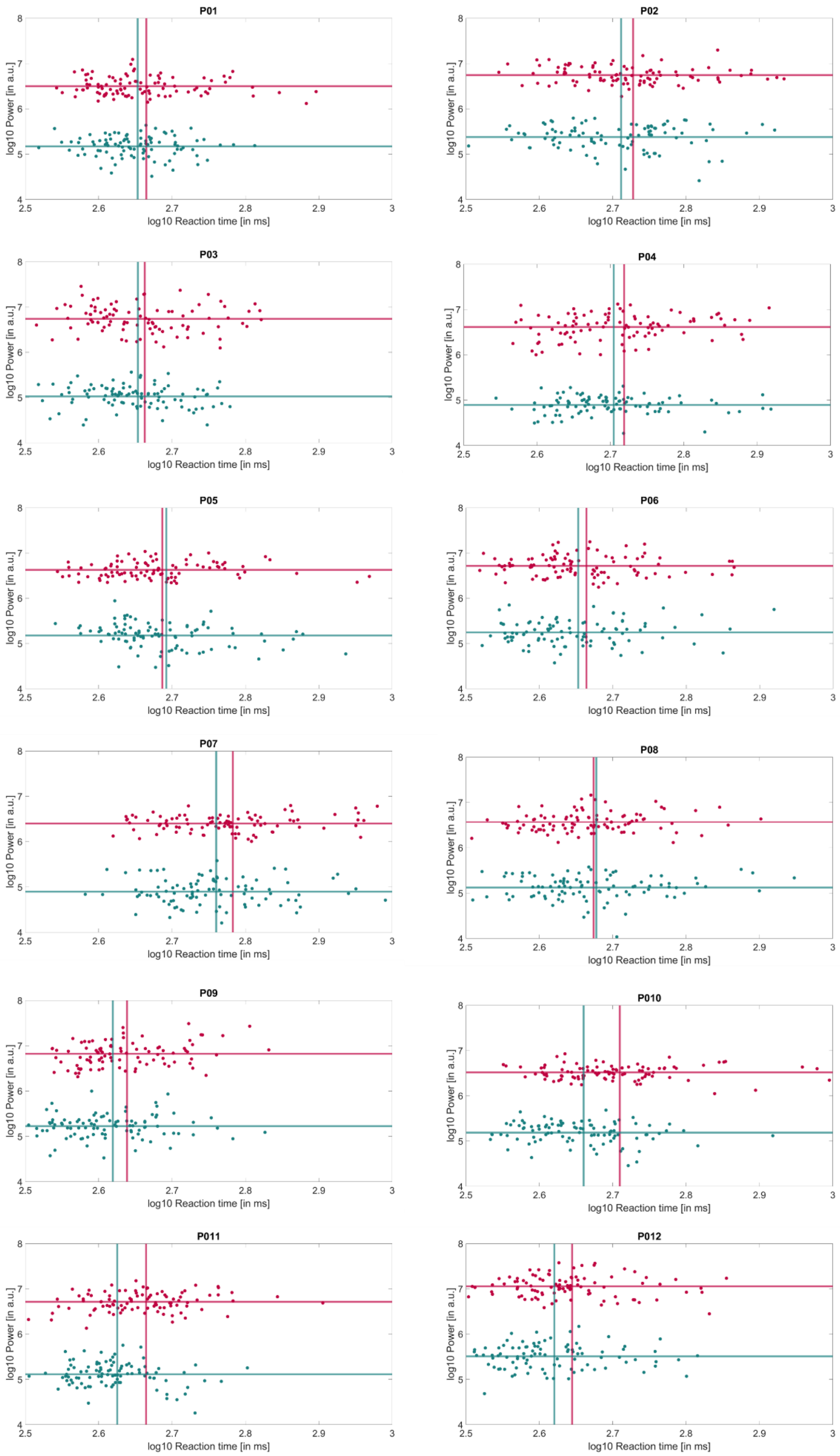


**SUPPLEMENTARY FIGURE 4. Individual relationship between the log-transformed mean power at the time-window of interest (TWoI) and the reaction time (RT) for burst (in red) and no-burst (in green) for validated trials.** Horizontal solid lines denote the mean power of burst (in red) and no-burst trials (in green), and vertical solid lines denote the mean RT of burst (in red) and no-burst trials (in green). Dots denote the mean power at the TWoI for each trial at individual level.

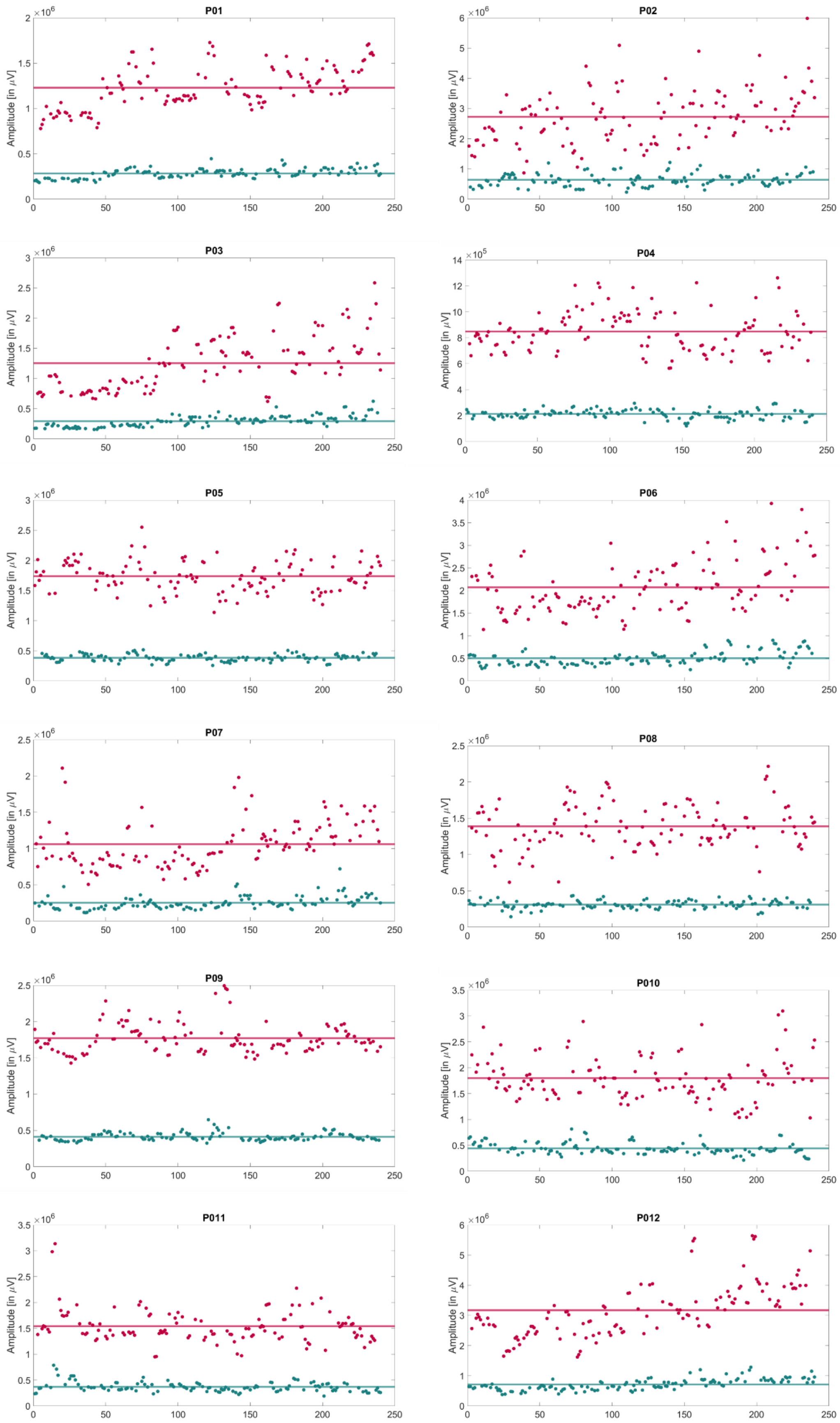


**SUPPLEMENTARY FIGURE 5. Individual power thresholds for burst (in red) and no-burst (in green) for validated trials.** Dots denote the amplitude thresholds for each trial at individual level for burst (in red) and no-burst trials (in green). Horizontal solid lines denote the mean power threshold for the cloud of dots of burst (in red) and no-burst trials (in green).

### REALITY CHECK: Selection of the frequency of interest (IFoI)

**SUPPLEMENTARY TABLE 3. Individual difference between amplitude [in dB] and frequency peak [in Hz] of the Individual Frequency of Interest (IFoI) during the task and during the training using Oz-electrode and OP-cluster of electrodes**. For each participant, the amplitude and frequency peak of the IFoI during the task and during the training for Oz-electrode and OP-cluster are provided, together with the difference in amplitude and frequency peak between task and training for Oz-electrode, and the difference in frequency peak between OP-cluster and Oz-electrode during the task and the training. NaN denotes that a peak was not found in the power spectrum (P02).

| Part. | Oz-electrode | | | | OP-cluster | | | | ΔIFoI Oz task – training | | ΔIFoI OP – Oz | |
|---|---|---|---|---|---|---|---|---|---|---|---|---|
| | IFoI (training) | | IFoI (task) | | IFoI (training) | | IFoI (task) | | | | Training | Task |
| | Peak [in Hz] | Amp. [in dB] | Peak [in Hz] | Amp. [in dB] | Peak [in Hz] | Amp. [in dB] | Peak [in Hz] | Amp. [in dB] | Peak [in Hz] | Amp. [in dB] | Peak [in Hz] | Peak [in Hz] |
| **1** | 10.25 | 5.70 | 10.00 | 6.87 | 10.25 | 3.75 | 10.00 | 2.22 | **0.25** | -1.17 | **0.00** | **0.00** |
| **2** | 10.75 | 2.97 | 10.25 | 2.19 | 10.50 | -10.50 | NaN | NaN | **-0.50** | -0.78 | **-0.25** | **NaN** |
| **3** | 10.50 | 9.91 | 10.50 | 9.11 | 10.50 | 8.55 | 10.50 | 7.91 | **0.00** | -0.80 | **0.00** | **0.00** |
| **4** | 10.50 | 11.26 | 10.25 | 10.85 | 10.50 | 11.24 | 10.50 | 10.82 | **-0.25** | -0.41 | **0.00** | **0.25** |
| **5** | 11.00 | 2.23 | 10.75 | 3.89 | 11.00 | -0.25 | 10.75 | -0.91 | **-0.25** | 1.65 | **0.00** | **0.00** |
| **6** | 10.00 | 2.00 | 9.75 | 4.29 | 9.50 | 1.54 | 9.50 | 2.62 | **-0.25** | 2.29 | **-0.50** | **-0.25** |
| **7** | 11.25 | 3.73 | 11.00 | 5.06 | 11.25 | 2.16 | 11.00 | 3.86 | **-0.25** | 1.33 | **0.00** | **0.00** |
| **8** | 11.25 | 5.36 | 11.00 | 2.68 | 11.25 | 3.90 | 11.00 | 3.16 | **-0.25** | -2.69 | **0.00** | **0.00** |
| **9** | 10.50 | 6.72 | 10.25 | 6.94 | 10.50 | 5.01 | 10.50 | 6.20 | **-0.25** | 0.22 | **0.00** | **0.25** |
| **10** | 10.25 | 2.18 | 10.00 | 2.79 | 10.25 | 0.55 | 10.00 | 1.11 | **-0.25** | 0.61 | **0.00** | **0.00** |
| **11** | 10.75 | 2.57 | 10.50 | 6.04 | 10.75 | 0.25 | 10.75 | 6.00 | **-0.25** | 3.47 | **0.00** | **0.25** |
| **12** | 10.25 | 6.78 | 10.00 | 7.97 | 10.25 | 6.90 | 10.25 | 8.21 | **-0.25** | 1.19 | **0.00** | **0.25** |
| **Mean (SD)** | **10.60 (0.41)** | **5.12 (3.11)** | **10.35 (0.41)** | **5.72 (2.73)** | **10.54 (0.49)** | **2.76 (5.46)** | **10.43 (0.46)** | **4.65 (3.49)** | **-0.25 (0.11)** | **0.61 (1.63)** | **-0.06 (0.16)** | **0.07 (0.16)** |

### EXPLORATORY ANALYSIS: RT fits using the ex-Gaussian function

**SUPPLEMENTARY TABLE 4. Individual ex-Gaussian fit parameters for burst and no-burst trials.** The mean (μ), standard deviation (σ), the exponential parameters (τ), and the log-likelihood (fVal) are provided for burst and no-burst trials. Individuals assess of statistical difference in the RT from the one-tailed permutation test ($p<.05$). 2 participants (P11, P12) showed a significance difference in the μ RT between burst and no-burst trials, 1 participant (P11) showed a significant difference in σ RT between burst and no-bursts, and 2 other participants (P01, P03) showed a significance difference in the τ RT between burst and no-burst trials.

| **Part.** | **Burst** | | | **No-burst** | | | **Diff. μ RT [in ms]** | **p** | **Diff. σ RT [in ms]** | **p** | **Diff. τ RT [in ms]** | **p** |
|---|---|---|---|---|---|---|---|---|---|---|---|---|
| | **μ RT [in ms]** | **σ RT [in ms]** | **τ RT [in ms]** | **μ RT [in ms]** | **σ RT [in ms]** | **τ RT [in ms]** | | | | | | |
| **1** | 381 | 20 | 82 | 409 | 43 | 41 | -28 | .99 | -23 | **.01** | 41 | **.007** |
| **2** | 438 | 53 | 97 | 439 | 68 | 76 | -1 | .52 | -15 | .35 | 21 | .40 |
| **3** | 382 | 36 | 78 | 410 | 50 | 41 | -28 | .94 | -15 | .25 | 38 | **.02** |
| **4** | 417 | 42 | 106 | 419 | 35 | 88 | -1 | .52 | 7 | .68 | 18 | .39 |
| **5** | 399 | 36 | 87 | 407 | 28 | 85 | -8 | .73 | 7 | .37 | 2 | .93 |
| **6** | 376 | 34 | 86 | 364 | 34 | 86 | 12 | .24 | 0 | .99 | -1 | .98 |
| **7** | 496 | 61 | 110 | 474 | 48 | 101 | 22 | .17 | 13 | **.01** | 9 | **.01** |
| **8** | 355 | 37 | 96 | 344 | 19 | 110 | 11 | .17 | 17 | .80 | -14 | .32 |
| **9** | 372 | 36 | 63 | 351 | 28 | 65 | 21 | .08 | 8 | .73 | -2 | .89 |
| **10** | 409 | 34 | 103 | 384 | 37 | 73 | 25 | .11 | -3 | .90 | 30 | .22 |
| **11** | 402 | 44 | 60 | 370 | 24 | 52 | 32 | **.02** | 20 | .08 | 8 | .63 |
| **12** | 370 | 37 | 71 | 344 | 28 | 73 | 26 | **.01** | 10 | .31 | -2 | .90 |
| **Mean (SD)** | **400 (38)** | **39 (10)** | **86 (17)** | **393 (41)** | **37 (14)** | **74 (22)** | **7 (20)** | **.13** | **2 (14)** | **.27** | **12 (17)** | **.01** |

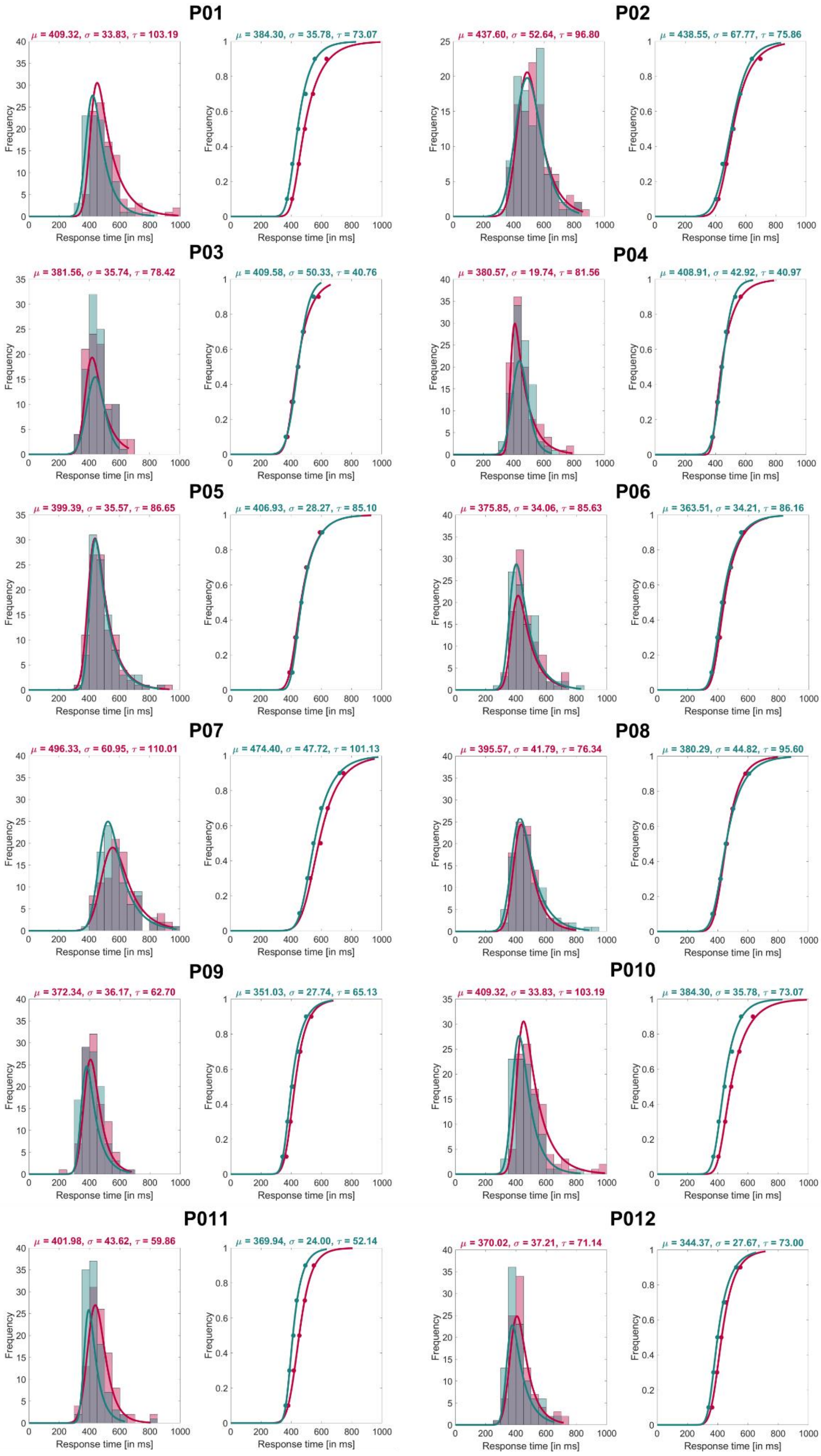


**SUPPLEMENTARY FIGURE 6. Individual ex-Gaussian fits for burst (in red) and no-burst (in green) trials.** For each participant, the histogram of burst and no-burst trials (on the left) is shown together with the probability density function curve of each trial condition (on the right).

### EXPLORATORY ANALYSIS: Commission and omission error rates for (no-)burst trials

**SUPPLEMENTARY TABLE 5. Individual error rates for go and no-go trials**. For each participant, the number of trials, the number of errors, and the error rate in each condition (go/burst, go/no-burst, no-go/burst, no-go/no-burst) are reported. Group-level t-tests assess statistical difference in error rates for burst and no-burst conditions for go and no-go trials.

| Part. | Go | | | | No-go | | | |
|---|---|---|---|---|---|---|---|---|
| | Burst | | No-burst | | Burst | | No-burst | |
| | No. of trials | Omission error rate | No. of trials | Omission error rate | No. of trials | Commission error rate | No. of trials | Commission error rate |
| 1 | 116 | **0.17** | 110 | 0.13 | 24 | **0.17** | 24 | 0.13 |
| 2 | 121 | **0.21** | 111 | 0.14 | 24 | 0.04 | 24 | **0.13** |
| 3 | 97 | **0.01** | 96 | 0.00 | 24 | 0.04 | 24 | **0.08** |
| 4 | 116 | 0.17 | 115 | 0.17 | 24 | **0.08** | 24 | 0.04 |
| 5 | 110 | **0.13** | 105 | 0.09 | 24 | 0.13 | 24 | 0.13 |
| 6 | 99 | 0.03 | 99 | 0.03 | 24 | 0.04 | 24 | **0.08** |
| 7 | 121 | 0.21 | 130 | **0.26** | 24 | 0.00 | 24 | **0.17** |
| 8 | 104 | **0.08** | 97 | 0.01 | 24 | 0.08 | 24 | **0.17** |
| 9 | 135 | **0.29** | 120 | 0.20 | 24 | 0.33 | 24 | **0.42** |
| 10 | 117 | **0.18** | 109 | 0.12 | 24 | **0.13** | 24 | 0.08 |
| 11 | 101 | **0.05** | 97 | 0.01 | 24 | 0.33 | 24 | **0.38** |
| 12 | 108 | **0.11** | 101 | 0.05 | 24 | 0.04 | 24 | **0.13** |
| **Mean (SD)** | **112 (11)** | **0.14 (0.08)** | **108 (10)** | **0.10 (0.08)** | **24 (0)** | **0.12 (0.11)** | **24 (0)** | **0.16 (0.12)** |

### EXPLORATORY ANALYSIS: Phase-behaviour opposition

**SUPPLEMENTARY TABLE 6. Individual phase opposition sum (POS) at stimulus onset for valid-burst trials between fast and slow RT.** For each participant, the mean (SD) RT [in ms] and phase [in degrees] of the overall trials, the number of trials, mean (SD) RT [in ms] and mean (SD) phases [in degrees] for fast and slow trials, the difference in RT [in ms] and in phases [in degrees] between fast and slow trials are provided. Individuals and group-level assess of statistical difference in phases from the phase opposition sum (POS) method.

| Part. | Mean (SD) RT [in ms] | Mean (SD) phase [in °] | FAST RT | | | SLOW RT | | | Diff. RT [in ms] | Diff. phases [in °] | POS | | |
|---|---|---|---|---|---|---|---|---|---|---|---|---|---|
| | | | No. trials | Mean (SD) RT [in ms] | Mean (SD) phase [in °] | No. trials | Mean (SD) RT [in ms] | Mean (SD) phase [in °] | | | POS value | Mean (SD) surrogate POS value | p |
| 1 | 462 (84) | 172 (77) | 48 | 404 (23) | -180 (79) | 48 | 521 (82) | 169 (75) | **117** | **-11** | <.001 | 0.061 (0.064) | .94 |
| 2 | 534 (107) | -178 (77) | 48 | 454 (43) | -165 (78) | 48 | 615 (91) | 176 (75) | **161** | **-18** | 0.002 | 0.052 (0.059) | .86 |
| 3 | 460 (80) | 89 (76) | 48 | 397 (30) | 56 (74) | 48 | 522 (63) | 128 (75) | **125** | **72** | 0.056 | 0.055 (0.062) | .36 |
| 4 | 523 (104) | -1 (78) | 48 | 442 (39) | -20 (78) | 48 | 604 (82) | 16 (77) | **162** | **35** | 0.007 | 0.076 (0.074) | .85 |
| 5 | 486 (99) | 141 (80) | 48 | 420 (33) | -172 (75) | 48 | 552 (99) | 25 (77) | **133** | **-157** | 0.192 | 0.125 (0.092) | .22 |
| 6 | 461 (90) | 154 (75) | 48 | 396 (31) | 143 (71) | 48 | 527 (80) | -169 (78) | **131** | **48** | 0.020 | 0.035 (0.043) | .48 |
| 7 | 606 (123) | -161 (76) | 48 | 516 (52) | -147 (71) | 48 | 697 (105) | 138 (78) | **182** | **-75** | 0.042 | 0.038 (0.048) | .32 |
| 8 | 472 (86) | -142 (77) | 48 | 408 (34) | -108 (75) | 48 | 536 (74) | -179 (76) | **128** | **-71** | 0.047 | 0.055 (0.061) | .41 |
| 9 | 435 (70) | 150 (76) | 48 | 382 (28) | 172 (72) | 48 | 488 (59) | 86 (77) | **106** | **-86** | 0.066 | 0.046 (0.055) | .26 |
| 10 | 512 (115) | 45 (74) | 48 | 435 (34) | -83 (74) | 48 | 589 (116) | 17 (71) | **154** | **-67** | 0.064 | 0.035 (0.046) | .18 |
| 11 | 462 (76) | -77 (76) | 48 | 406 (32) | -27 (74) | 48 | 518 (65) | -122 (74) | **112** | **-95** | 0.106 | 0.060 (0.067) | .20 |
| 12 | 441 (82) | 178 (75) | 48 | 384 (32) | -150 (77) | 48 | 499 (76) | 163 (72) | **115** | **-47** | 0.022 | 0.031 (0.041) | .42 |
| **Mean (SD)** | **488 (49)** | **-** | **48** | **420 (38)** | **-** | **48** | **556 (60)** | **-** | **135 (24)** | **-43 (57)** | **0.052** | **0.056 (0.066)** | **.37** |

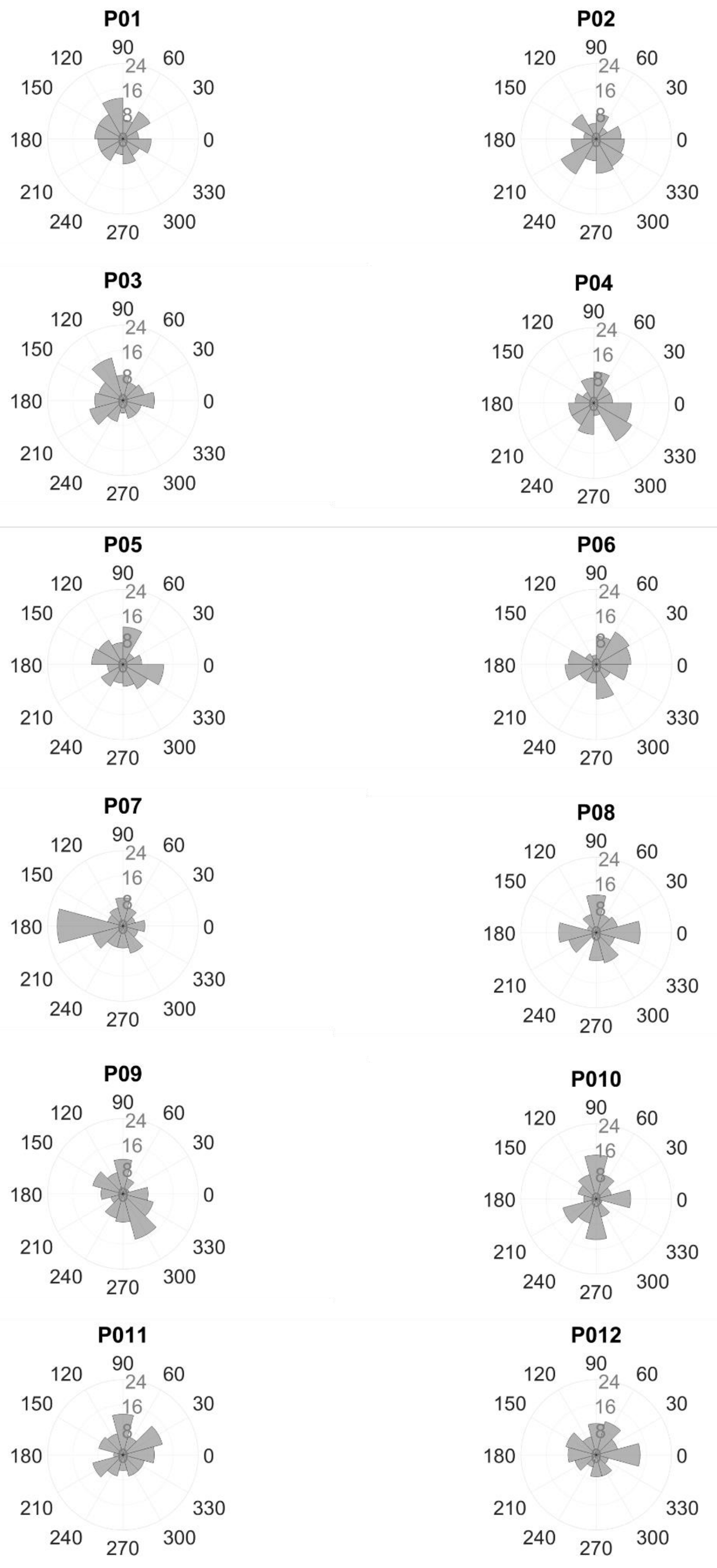


**SUPPLEMENTARY FIGURE 7. Individual rose plot of the phase distribution at stimulus onset for valid-burst trials [in degrees].** Individual rose plot of phases for all trials for each participant. Each bin corresponds to 30º.

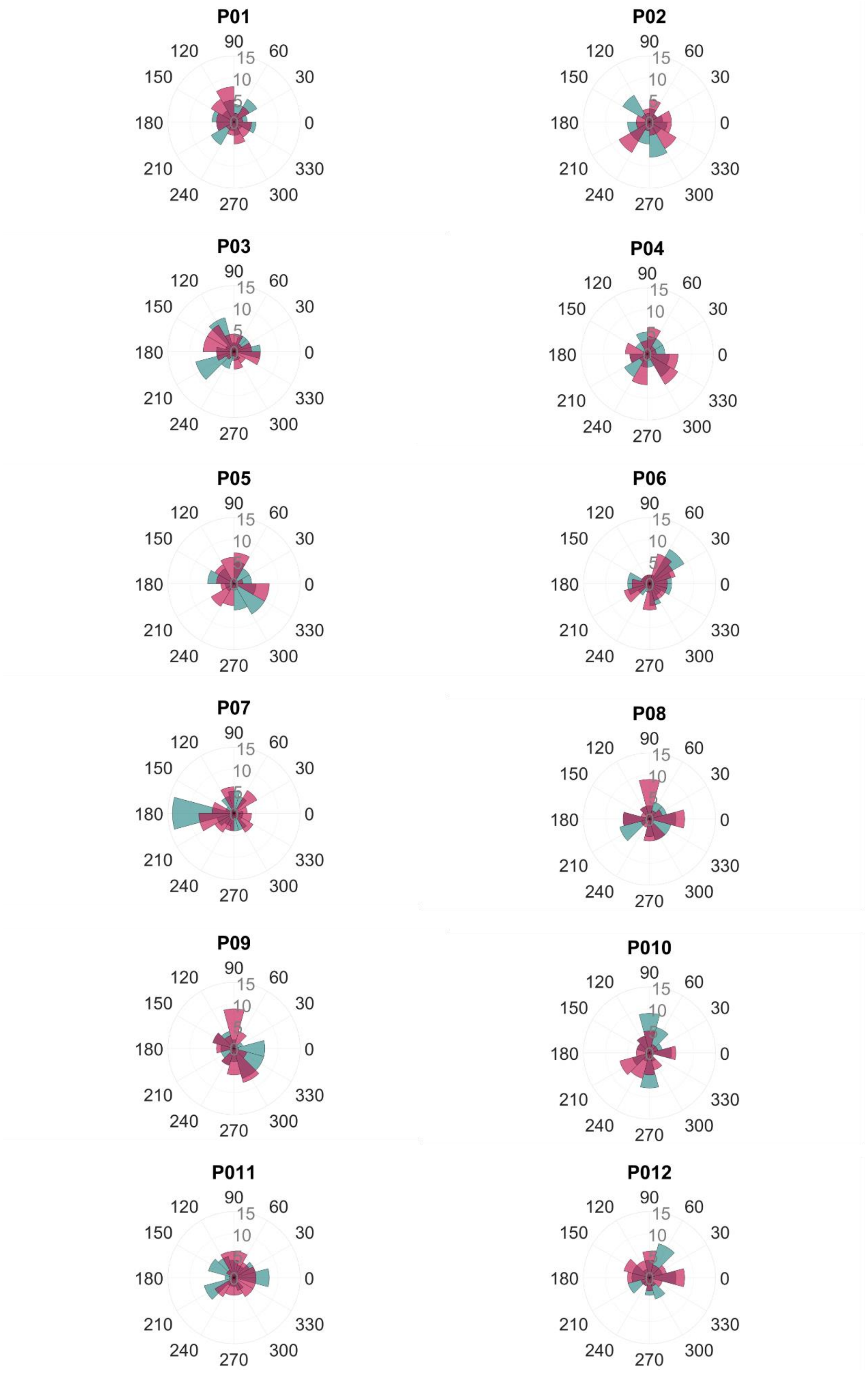


**SUPPLEMENTARY FIGURE 8. Individual rose plot of the phase distribution at stimulus onset for valid-burst trials [in degrees].** Individual rose plot of phases for fast (in green) and slow (in red) trials for each participant. Each bin corresponds to 30º.

### EXPLORATORY ANALYSIS: Exclusion of trials due to power criterion

In order to achieve the intended number of valid trials (n=240), we collected an average of 654 (SD = 79) responses per participant before filtering online according to preregistered values (see *Methods* section). On average per participant, 28 (SD = 21; 4%) trials were excluded for not satisfying the *reaction time criterion*, 344 (SD = 78; 52%) for not meeting the *power threshold* criterion, and 43 (SD = 25; 7%) because of the *artefact* criterion (see **Supplementary Table 7** for more details). Note that the exclusion of trials proceeded in the sequential order artefact, then power, then RT criterion, and once a trial was dropped, no further checks were done on the remaining criteria.

**SUPPLEMENTARY TABLE 7. Individual number of excluded trials.** For each participant, it is reported the total number of trials delivered, the sum of all the trials excluded for not satisfying the criteria of the stud (including the percentage; %), the number of trials excluded for reaction time (RTs), the number of trials excluded for not satisfying the power threshold criterion, and the number of trials excluded for not satisfying the artifact threshold criterion. The difference between the number of all trials and the number of excluded trials is the number of valid trials (N = 240).

| Part. | No. all trials | All | | RT | | Power | | Artifact | |
|---|---|---|---|---|---|---|---|---|---|
| | | No. excl. trials | % excl. trials | No. excl. trials | % excl. trials | No. excl. trials | % excl. trials | No. excl. trials | % excl. trials |
| **1** | **760** | **520** | 68% | **34** | 4% | **462** | 61% | **24** | 3% |
| **2** | **738** | **498** | 67% | **40** | 5% | **361** | 49% | **97** | 13% |
| **3** | **580** | **340** | 59% | **1** | 0% | **308** | 53% | **31** | 5% |
| **4** | **574** | **334** | 58% | **39** | 7% | **272** | 47% | **23** | 4% |
| **5** | **616** | **376** | 61% | **23** | 4% | **318** | 52% | **35** | 6% |
| **6** | **588** | **348** | 59% | **6** | 1% | **298** | 51% | **44** | 7% |
| **7** | **571** | **331** | 58% | **59** | 10% | **227** | 40% | **45** | 8% |
| **8** | **669** | **429** | 64% | **9** | 1% | **404** | 60% | **16** | 2% |
| **9** | **725** | **485** | 67% | **63** | 9% | **372** | 51% | **50** | 7% |
| **10** | **724** | **484** | 67% | **34** | 5% | **375** | 52% | **75** | 10% |
| **11** | **565** | **325** | 58% | **6** | 1% | **259** | 46% | **60** | 11% |
| **12** | **741** | **501** | 68% | **17** | 2% | **473** | 64% | **11** | 1% |
| **Mean (SD)** | **654 (79)** | **414 (79)** | ***63% (4%)*** | **28 (21)** | **4% (3%)** | **344 (78)** | **52% (7%)** | **43 (25)** | **7% (4%)** |

We decided to pay a closer look to the average of 52% (SD = 7%) of trials (**Supplementary Table 7**) excluded for not satisfying the *(no-)burst power threshold* criterion (see *Trial validity criteria*). To further explore the data, we selected the trials excluded for power criteria (which satisfied RT and artifact power criteria) and divided them into burst and no-burst categories. The mean number of trials excluded for power in burst trials was 116 (SD = 63; Max = 251; Min = 46), whereas in no-burst trials the mean number was 220 (SD = 77; Max = 404; Min = 124) (**Supplementary Table 8**). Overall, 10 out of twelve participants excluded more trials in no-burst trials compared to burst trials. These results show that it was more difficult to detect non-oscillatory activity during the task execution in comparison of detecting oscillatory burst activity in the EEG signal for most of the participants.

**SUPPLEMENTARY TABLE 8. Individual IFoI amplitude, the number of excluded trials for burst and no-burst trials, and the mean (SD) percentage of data points above/below the threshold for burst and no-burst trials, respectively.** For each participant, the amplitude [in dB] of the Individual Frequency of Interest (IFoI) at task using Oz-electrode is given with the number of trials excluded for power, and for burst and no-burst trials separately.

| Part. | IFoI (task) amp. [in dB] | No. excl. trials for power | No. burst trials | No. no-burst trials |
|---|---|---|---|---|
| **1** | 5.70 | 462 | 97 | **365** |
| **2** | 2.97 | 361 | **204** | 157 |
| **3** | 9.91 | 308 | 46 | **262** |
| **4** | 11.26 | 272 | 61 | **211** |
| **5** | 2.23 | 318 | 130 | **188** |
| **6** | 2.00 | 298 | 140 | **158** |
| **7** | 3.73 | 227 | 72 | **155** |
| **8** | 5.36 | 404 | 155 | **249** |
| **9** | 6.72 | 372 | 92 | **280** |
| **10** | 2.18 | 375 | **251** | 124 |
| **11** | 2.57 | 259 | 72 | **187** |
| **12** | 6.78 | 473 | 69 | **404** |
| **Mean (SD)** | **5.12 (3.11)** | **344 (78)** | **116 (63)** | **220 (77)** |
